# Single-molecule, full-length transcript isoform sequencing reveals disease mutation-associated RNA isoforms in cardiomyocytes

**DOI:** 10.1101/2021.06.18.448952

**Authors:** Chenchen Zhu, Jingyan Wu, Han Sun, Francesca Briganti, Benjamin Meder, Wu Wei, Lars M. Steinmetz

## Abstract

Alternative splicing generates differing RNA isoforms that govern phenotypic complexity of eukaryotes. Its malfunction underlies many diseases, including cancer and cardiovascular diseases. Comparative analysis of RNA isoforms at the genome-wide scale has been difficult. Here, we established an experimental and computational pipeline that performs *de novo* transcript annotation and accurately quantifies transcript isoforms from cDNA sequences with a full-length isoform detection accuracy of 97.6%. We generated a searchable, quantitative human transcriptome annotation with 31,025 known and 5,740 novel transcript isoforms (http://steinmetzlab.embl.de/iBrowser/). By analyzing the isoforms in the presence of RNA Binding Motif Protein 20 (*RBM20*) mutations associated with aggressive dilated cardiomyopathy (DCM), we identified 121 differentially expressed transcript isoforms in 107 cardiac genes. Our approach enables quantitative dissection of complex transcript architecture instead of mere identification of inclusion or exclusion of individual exons, as exemplified by novel IMMT isoforms. Thereby we achieve a path to direct differential expression testing independent of an existing annotation of transcript isoforms as the functional unit, instead of genes or exons, providing more immediate biological interpretation and higher resolution transcriptome comparisons.

---

Nearly all human multi-exon genes are alternatively spliced^1-4^, allowing a single gene to generate multiple RNA isoforms that give rise to different protein isoforms^5,6^, which consequently drive phenotypic complexity in eukaryotes^1^. Numerous diseases including cancers, neurological and cardiovascular diseases, are associated with alternative splicing dysregulation, such as mutations in the heart-specific alternative splicing regulator, *RBM20*, that cause dilated cardiomyopathy (DCM)^7-9^. The conventional method to study alternative splicing is short-read RNA-seq^1,4,10-13^, which sequences fragments of RNA and can therefore only detect expression changes of the whole gene or individual exons. Although software tools such as Kallisto^14^ or Salmon^15^ permit transcript level quantification with short-read RNA-seq, this requires either a reference annotation or de novo transcriptome assembly. The latter, needed for discovery of novel isoforms, is particularly challenging from short read data. Therefore, short-read RNA-seq is not ideal for quantitatively detecting individual RNA isoforms. The emergence of long-read sequencing technologies including Pacific Biosciences (PacBio) and Oxford Nanopore Technologies (ONT) have provided a new tool for more comprehensive analysis of alternative splicing. An initial, genome-wide study with PacBio used 476,000 reads^16^, which enables *de novo* transcript identification but is insufficient for accurate quantification. Consequently, subsequent studies with PacBio or ONT focused on quantifying selected genes using enrichment methods, lacking genome-wide profiles^17-20^. Recent studies have quantified transcripts using long-read sequencing in mouse^21,22^ and breast cancer cells^23^, however no quantitative differences between samples were assessed. Furthermore, long-read sequencing has just begun to be utilized in single-cell transcriptomics for studying transcriptional heterogeneity^24,25^. It is challenging to apply long-read RNA-seq to determine global expression changes of transcripts, because: 1) low sequencing depth often doesn’t allow adequate genome coverage for accurate quantification; 2) high sequencing error rates require novel computational methods that allow transcript quantification, classification and visualization; 3) high false discovery rates of identified unannotated transcripts due to sequencing artifacts such as undesired template switching of reverse transcriptase (RT) and off-target priming of oligo(dT)^21^ confound downstream analysis. Thus, quantitative, genome-wide differential expression analysis to detect isoform changes at scale has not been possible so far. This has made it difficult to identify the molecular mechanisms underlying many diseases in which splicing has been implicated.

Here, we generated a large, long-read dataset for human induced pluripotent stem cell derived cardiomyocytes (iPSC-CMs) using ONT sequencing of cDNA and developed a workflow that accurately and quantitatively measures and compares full-length splicing isoforms at genome-scale independent of an existing annotation. RNAs from human iPSC-CMs (each with two independent clones) with and without DCM-associated *RBM20* mutations, together with spike-in controls (ERCC^26^ and sequins^27^), were converted to full-length cDNAs and sequenced with ONT MinION technology (Method, Supplementary Fig. 1, and Supplementary Table 1). In total, we generated 21 million high quality reads (mean qscore>=6, Method, Supplementary Fig. 2), and quantified 36,765 transcript isoforms for a total of 11,707 genes, covering 53.8% (10,682/19,847) of the protein-coding genes in the human genome (Figure 1, Supplementary Fig. 3, Supplementary Table 1 - 3). In comparison, our complementary short-read data with about 1,200 million pair-end reads identified 16,726 protein-coding genes (read count > 1).

**Figure 1.**
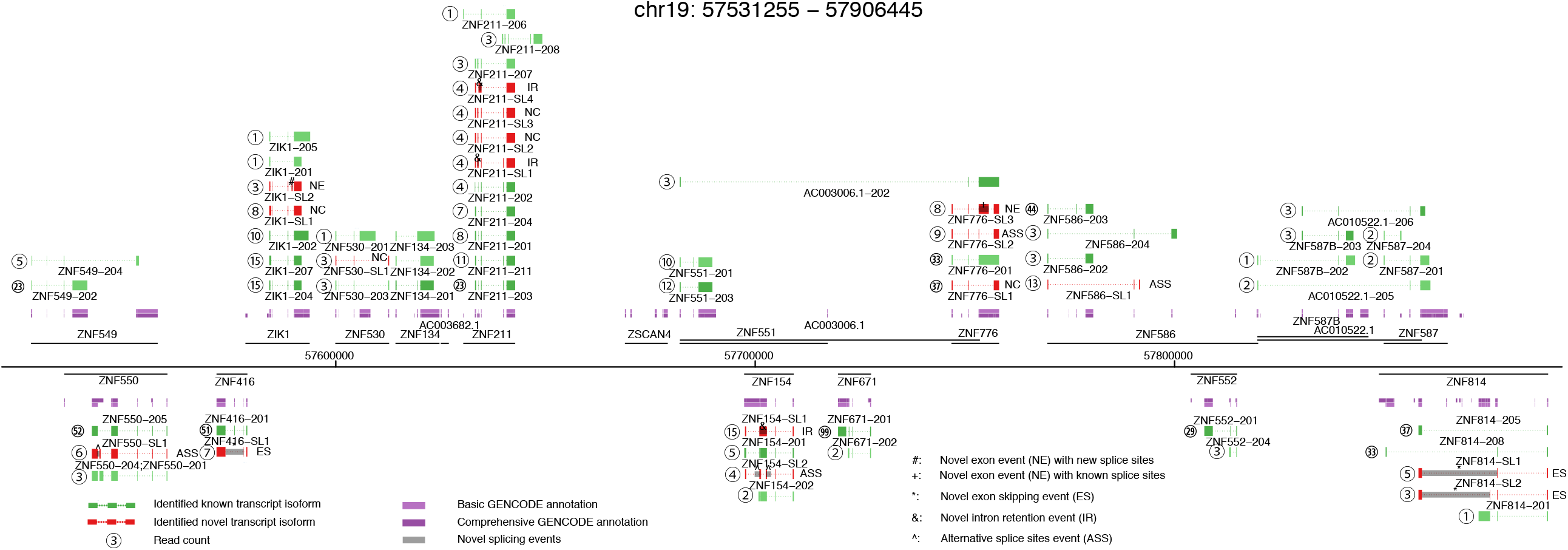
Complex landscape of full-length human iPSC-CM transcriptome. Genome-wide measurement of full-length splicing isoforms in human iPSC-CMs with a region of chromosome 19 (chr19:57531255-57906445) as an example. Gene loci are presented as a horizontal black line with annotated genes on + or – strands above or below the genome axis, respectively. Collapsed GENCODE comprehensive and basic annotation are presented in dark purple and purple track. Known transcript isoforms are shown as green tracks, and previously unidentified transcript isoforms as red. Known transcript isoforms didn’t pass transcript filtering criteria are presented in light green. Read counts for each identified transcript are given as number in circle. Based on the novel splicing events (location indicated with grey box and text symbols), novel transcript isoforms are categorized as novel exon combination (NC), novel exon (NE), novel intron retention (IR), novel exon skipping (ES), or novel alternative splice sites (ASS). Dubious novel read through transcripts are not shown.

Major problems of current methods to quantitate isoforms using long-read sequencing are sequencing artifacts and high sequencing errors resulting in high false transcript discoveries. To address these challenges, we established a computational method for full-length transcript quantification (FulQuant), which integrates accurate full-length read identification, transcript quantification and visualization from ONT cDNA sequences. This new method allows de novo transcript annotation and employs stringent and complex criteria for filtering reads, alignments, and transcripts, to rule out artifacts from RT and sequencing errors, and groups reads into transcripts based on their splice sites, generating a set of highly confident transcripts with abundance estimates (Methods, Supplementary Fig. 4). To evaluate FulQuant’s accuracy to identify isoforms, we determined its performance on synthetic sequencing spike-in sequences (sequins, spiked in at 3% of total RNA) spanning a ∼10^6^-fold range in concentration that covers the dynamic range of gene expression observed across the human transcriptome^27^. These spike-ins represent 72 artificial gene loci with multiple exons encoding 156 alternative isoforms to mimic human splice isoforms of different sequences, structures, abundances and lengths. We identified 82 of the 156 sequin isoforms, including 90% (70) of the 50% most concentrated isoforms, which account for 99.95% of all sequin RNA molecules. Our true positive rate for isoform identification, defined as the ratio of identified reference transcripts over all reported transcripts, was 97.6% based on all sequin isoforms, with the largest sequin isoform being ∼7 kb (Figure 2a). 98.1% of all spliced transcripts in human GENCODE annotation are below this 7 kb threshold (Figure 2b). The true positive rate was 100% for isoforms shorter than 1,728 bp, which is greater in length than 70% of all human spliced transcripts. The identification accuracy of FulQuant was higher than the existing long-read pipeline FLAIR^28^, which yielded a 30% true positive rate when benchmarked against our sequin dataset using default settings (Supplementary Text). FulQuant’s improvement in transcript identification over FLAIR comes primarily from removal of false positive transcripts resulting from artefacts in RT and sequencing that manifested in the synthetic controls. With FulQuant, the quantification of sequins correlated linearly with input concentrations (average Pearson correlation coefficient 0.85, Supplementary Fig. 5). Furthermore, 98.8% of transcript boundaries were positioned within 20 bp of the annotated transcripts (Supplementary Fig. 6), indicating a reliable capture of full-length transcripts without relying on annotation. Likewise, when we applied FulQuant to our human iPSC-CM dataset, we observed a high correlation between technical and biological replicates (Supplementary Fig. 7), and accurate boundary definition of full-length transcripts (Supplementary Table 2 and 3). FulQuant’s quantification generated better agreement between biological replicates than FLAIR (Supplementary Fig. 7), likely due to reduced number of false positive transcripts.

**Figure 2.**
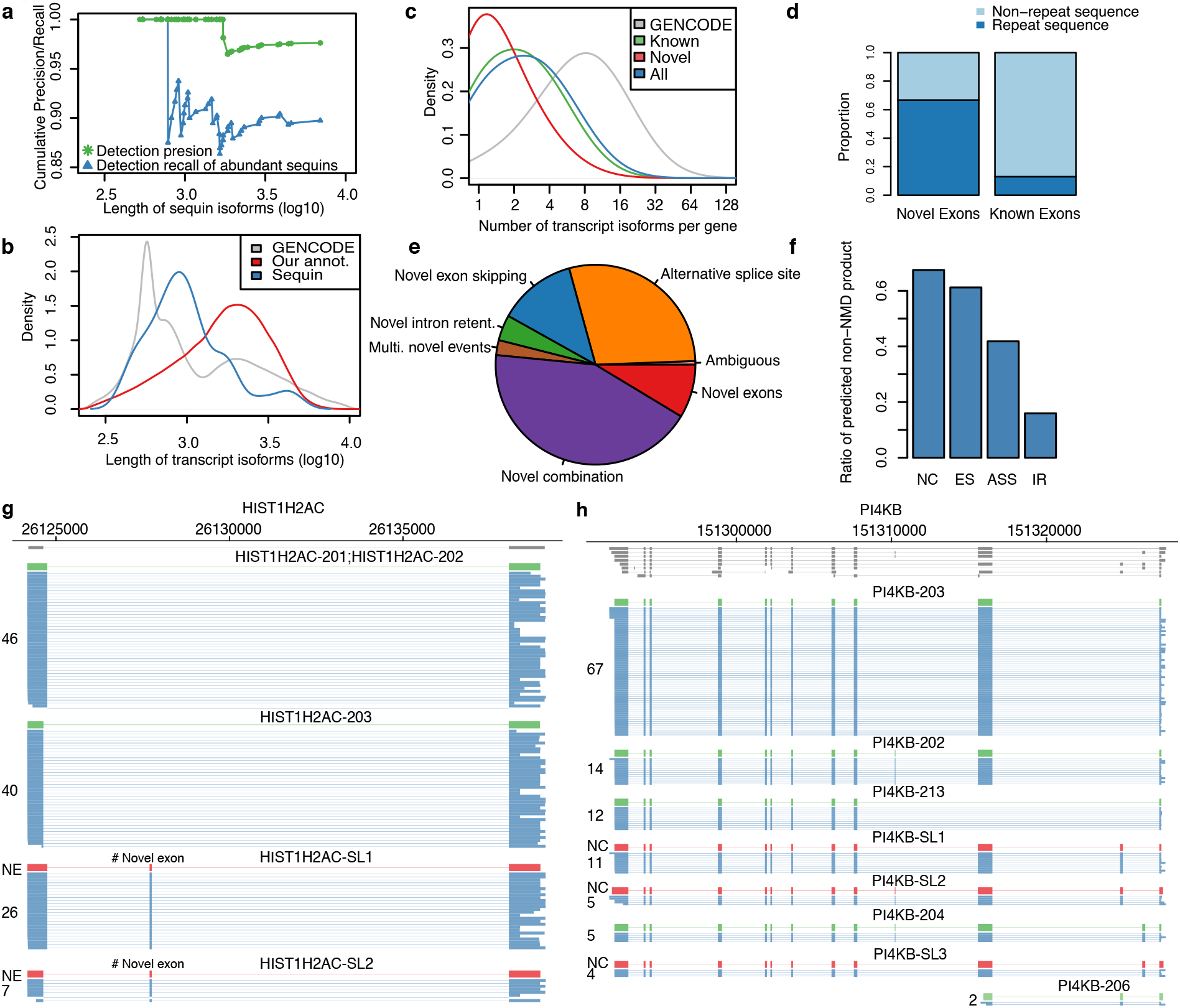
FulQuant method accurately identifies transcript isoforms in iPSC-CMs. (**a**) Length dependent precision (green) and recall (blue) of the FulQuant method as estimated using all or the most abundant 50% synthetic sequin controls. (**b**) Length distribution of transcripts in GENCODE, of our identified isoforms, and of the sequin controls. (a) and (b) are vertically aligned with the same x axis to illustrate the relevance of the accuracy estimated by sequins on human transcripts. (**c**) Distribution of number of isoforms per gene based on GENCODE comprehensive annotation and the transcripts identified in this study (divided into Known, Novel, and All). (**d**) Proportion of known exons that contain repeat sequences compared to the proportion of identified novel exons. (**e**) Percentages of novel transcripts isoforms by the types of novel splicing events: novel exon (NE), novel intron retention (IR), novel alternative splice sites (ASS), novel exon skipping (ES) and unannotated combinations of known exons (NC). Ambiguous transcripts are those with alignment issues. (**f**) Ratio of predicted non-NMD product in each novel transcript categories. (**g**) and (**h**) Examples of genes with unannotated combinations of known exons and novel exon, respectively. Known isoforms in the comprehensive GENCODE annotation are shown in gray track. Isoforms with less than 5 reads are not shown.

Our data revealed genome-wide complex alternative splicing patterns for known and novel transcript isoforms. By comparing our transcript isoforms to the human GENCODE annotation (V24), we identified 36,765 transcript isoforms, 5,740 (15.6%) of which were novel. For example, we identified eight known and four novel transcripts for the zinc-finger gene, *ZNF211*(Fig. 1). When comparing to a comprehensive annotation set CHESS (v2.2)^29^, 3028 (8.2%) were novel (Supplementary Text and Fig. 8). Our genome-wide full-length isoform dataset is available via a searchable browser, which we have developed and made freely available (http://steinmetzlab.embl.de/iBrowser/).

Our identified transcript isoforms have lengths from 177 bp to 10,718 bp with a median of 1,768 bp (Figure 2b and Supplementary Fig. 9). The length distribution is similar to GENCODE full-length protein-coding and lincRNA transcripts (Supplementary Fig. 10), however we didn’t capture large transcripts such as *TTN* isoforms (∼100 kb in size, a major structure component of the heart muscle) likely due to RT limitation of cDNA synthesis. A median of two alternatively spliced transcript isoforms per gene were identified (Figure 2c). It is notable that by applying our approach to only one cell type, namely iPSC-CMs, we identified already 24.1% of protein-coding isoforms (n=80,901) covering 9,946 genes in the GENCODE comprehensive annotation, which cumulates data from various human cell lines, tissues, and population. This reflects the comprehensive nature and depth of our long-read approach, and suggests that the number and complexity of all splice isoforms in the human genome are likely underestimated.

Aberrant splicing may generate novel transcripts, which could be used for diagnostics and drug development. Notably novel exons detected in our dataset were enriched in repetitive sequences defined by RepeatMasker (Figure 2d), with 25% of novel exons overlapping SINE/Alu sequences. In addition to a higher degree of intron retention (IR, 4.1%), most novel transcript isoforms contained alternative splice sites (ASS, 29.2%), novel exon skipping (ES, 12.7%), novel exons (NE, 8.6%) and unannotated combinations of known exons (NC, 42.9%, Figure 2e, g and h). To assess the performance of our pipeline, we have successfully validated 26 out of 33 novel splice events (exon skipping, intron retention and novel exons) using independent fragment analysis by RT-PCR (Methods, Supplementary Fig. 11 and 12 and Table. 6). In terms of protein-coding potential, our identified novel transcript isoforms add 1 potential protein-coding isoform every four genes on average (Figure 2f, Supplementary text). Supporting the resolution of our data, we also identified 45 potential spliced polycistronic transcripts from adjacent genes (Supplementary Fig. 13) and unspliced polycistronic transcripts that are transcribed from the mitochondrial genome (Supplementary Fig. 14)^30,31^.

Coordinated expression of exons underlies many regulatory mechanisms^23^. Its analysis also provides important insights into strategies of alternative splicing^32,33^. Particularly in heart, mutually exclusive exon pairs in genes such as TPM1, 2 and 3 have been used as model systems to study splicing regulation^34^. Our long-read data enabled a first systematic analysis of the interdependency between adjacent and distant exons across an entire transcript isoform in heart cells. We identified 2,793 significantly co-regulated exon pairs in 722 genes (adjusted P<0.001, mutually exclusive and inclusive association, Methods and Supplementary Table 4). For example, we detected the well-documented exon pair association in the heart-specific gene, TPM1, that harbors adjacent mutually exclusive exons 2 and 3, which are representative of muscle type switching between smooth and skeletal muscle^35^. In addition, we identified 6 further co-association events in TPM1 transcript isoforms, including 4 distal pairs, such as a mutually inclusive pair between exons 2 and 11 and a mutually exclusive pair between exons 7 and 11 (Figure 3a). Another prominent example is MYL7 (Figure 3b), a critical gene for cardiac function, that contains 18 mutually inclusive (6) and exclusive (12) exon pairs. Over 70% of all tested exon pairs (25) in this gene were significantly co-associated, and the majority (61%) are distally positioned, suggesting complex splicing regulation for this gene. Overall, we observed that adjacent co-association, within one or two exons, is more common than distal co-association (Supplementary Fig. 15). Since splicing of nascent RNAs occurs co-transcriptionally at an extremely fast rate (“50% of splicing is complete within ∼1.4 s after 3′ splice site synthesis”)^36^, we reasoned that this theoretically favors prompt correlated splicing of adjacent, newly synthesized exons.

**Figure 3.**
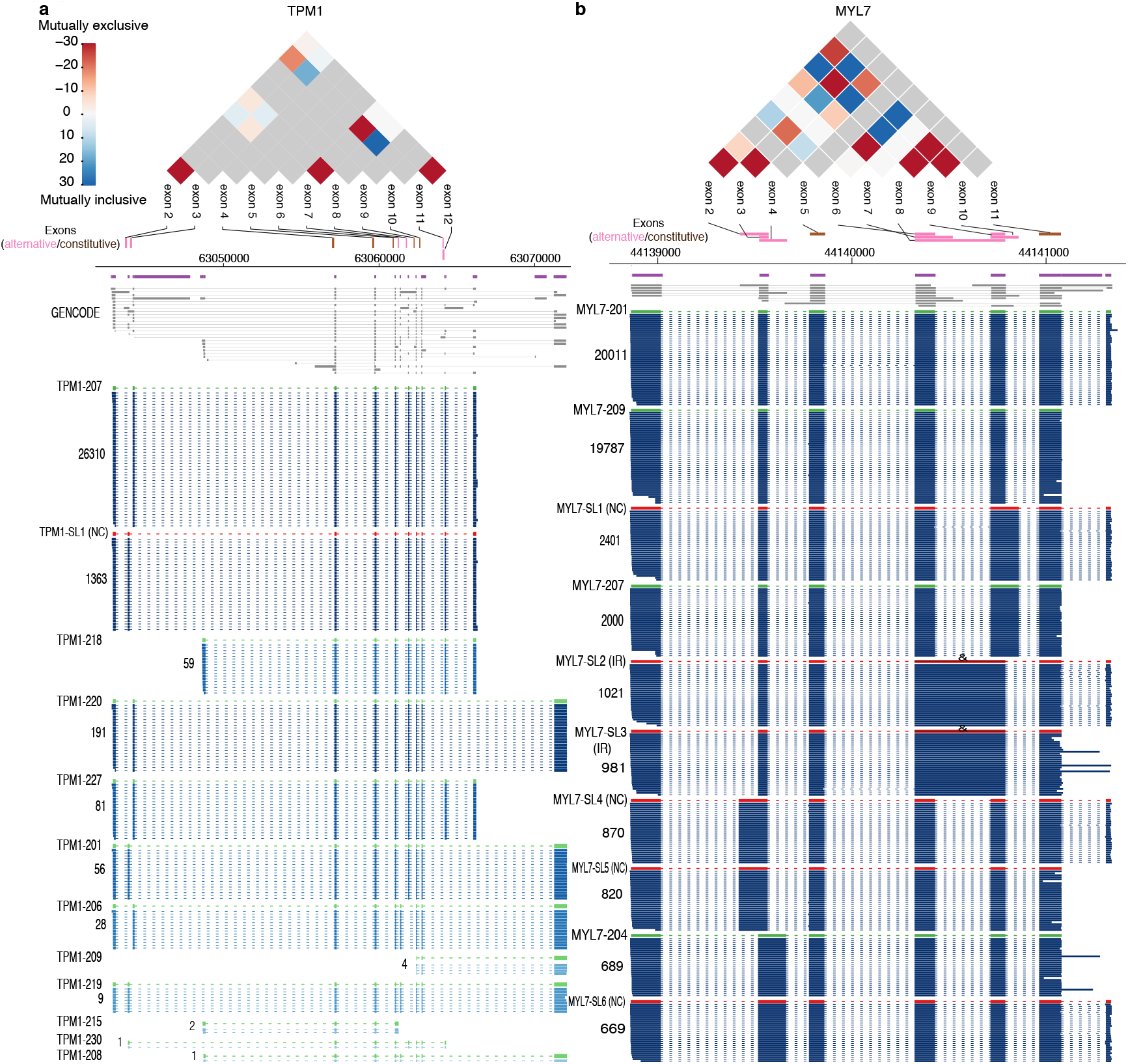
Exon co-associated events in human iPSC-CMs. TPM1 (**a**) and MLY7 (**b**) are shown as examples of genes containing multiple adjacent and distal mutually inclusive or exclusive alternative exons. The co-association strength (log10 of p-value) between alternative exon pairs is illustrated in the triangle plot for mutually exclusive (red, -) and inclusive pairs (blue, +), while the gray color indicates exon pairs not tested. Purple and gray annotations are for collapsed and expanded GENCODE comprehensive annotation, respectively. Reads in blue are in the same log-scale as Figure 1 with darkness correlating with the read counts. Only exons used for our co-association analysis are shown. Lowly abundant transcript isoforms are not shown for illustration clarity.

Aberrant splicing is linked to a wide variety of diseases, however identifying the causative event is challenging. Our approach using full-length transcript sequencing provides an opportunity to study differential isoform expression between different states, which will enable the direct comparison between heath and disease. To develop a quantitative approach to identify differentially expressed transcript isoforms between sample types, we applied our pipeline to studying how full-length splicing isoforms are affected by mutations in the heart-specific splicing regulator *RBM20*^37-39^. *RMB20* mutations (such as R634Q within its arginine/serine-rich domain) lead to an aggressive form of DCM and short read sequencing has shown that *RBM20* target exons are mis-spliced in rat^37^, mouse^40^, pig^41^, and human cardiac cells^38,42^. However, in many of these targets it has remained unclear to which aberrantly spliced transcript isoform these differentially spliced exons belong and whether that may or may not produce malfunctioning proteins, or may simply lead to the absence of correctly spliced protein. In addition to the 6M reads for the R634Q mutant, we also generated 9M reads for the P633L mutant, a new pathogenic mutation (proline-to-leucine change at amino acid position 633) that we recently identified^42,43^. We performed *de novo* transcript annotation using R634Q and wildtype data. While this approach would not identify novel isoforms seen only in the P633L mutant, it allowed us to quantitatively compare the expression level of isoforms detected in R634Q and wild type, across all mutants. To compare full-length transcriptomes of iPSC-CMs harboring mutants R634Q and P633L to wild-type *RBM20*, we performed differential isoform expression analysis using DESeq2^44^ on the quantified transcripts (Supplementary Table 3). We identified 121 transcript isoforms in 107 genes that are differentially expressed (adjusted P<0.001, Figure 4a, Supplementary Table 5, Methods). Both mutants showed similar expression levels (correlation coefficient = 0.77) and didn’t differ significantly when compared to each other using differential expression analysis (Supplementary Fig. 16). Over 80% of our candidates identified by long-read sequencing were recapitulated by the same analysis run on the short-read data with transcript isoform quantification using Kallisto (Methods, Supplementary text and Fig. 17 and 18). Gene ontology analysis of these 107 genes showed enrichment in cardiac functions, like striated muscle development, regulation of heart contraction, ion transport, and actin filament-based processes (adjusted P<0.05), supporting the essential role of *RBM20* in cardiac function (Supplementary text and Supplementary Fig. 19).

**Figure 4.**
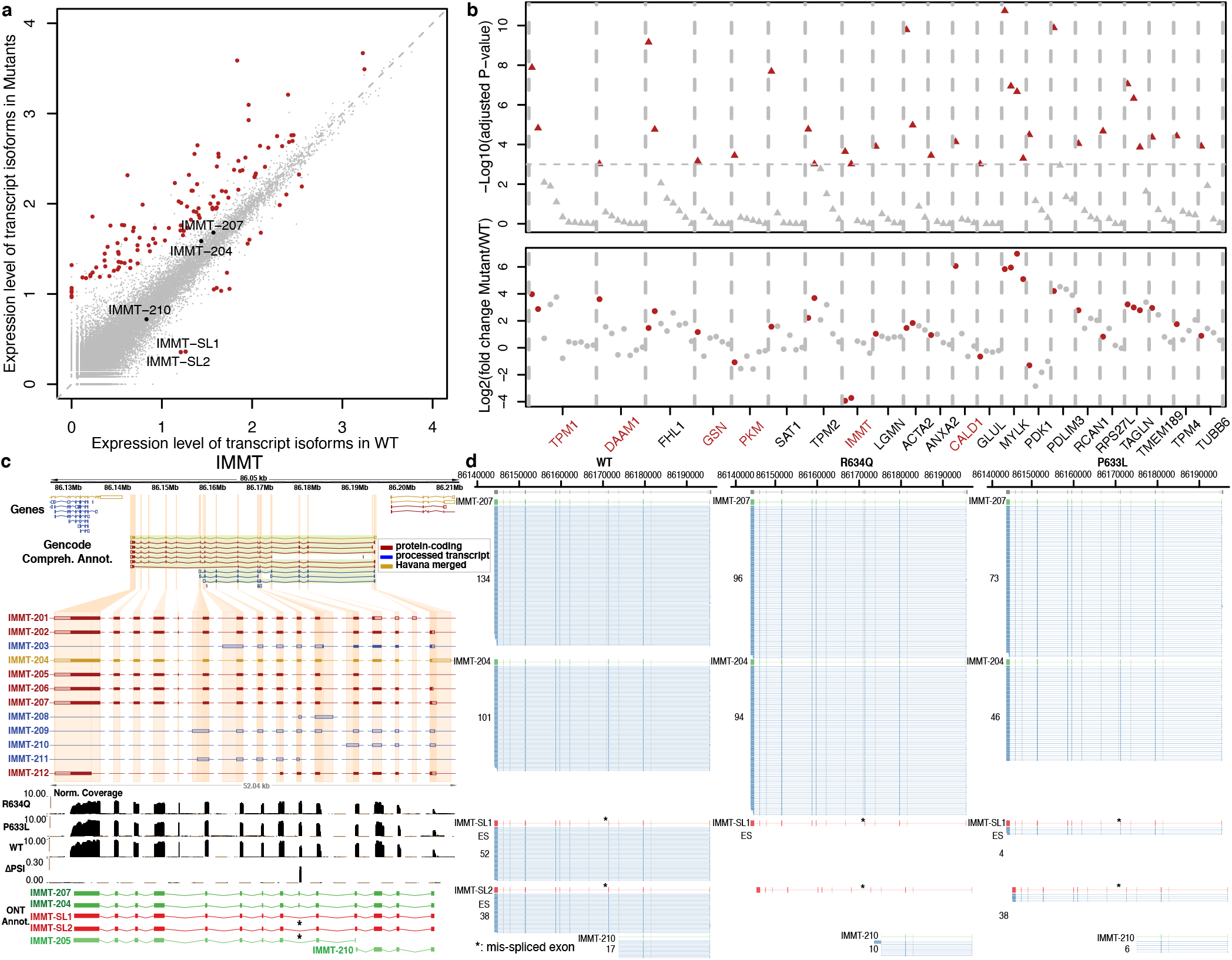
Differentially expressed full-length transcript isoforms in the *RBM20* mutants R643Q and P633L. (**a**) Scatter plot for average full-length isoform expression levels in WT and mutants. Each dot represents an isoform, with red dots highlighting the differentially expressed ones (adjusted P<0.001). Isoforms for *IMMT* gene are labeled. (**b**) Genes containing differentially expressed isoforms. Top, significance level of the differential expression analysis for each isoform (triangle) with significant ones in red. Bottom, fold change of expression level of each isoform (point) with significant ones in red. Gene names in red are genes differentially expressed only proportion of their isoforms, with no significant change at the total gene expression level. Only genes with more than three expressed isoforms are displayed. (**c**) *IMMT* gene with GENCODE annotation, short-read data and identified transcript isoforms in long-read data of this study. Tracks from top to bottom: collapsed view of GENCODE transcripts around *IMMT* locus, expanded view of *IMMT* GENCODE transcripts, short-read coverage for mutant and WT, delta-PSI between mutant and WT inferred in short-read data, and known (green) and novel (red) transcripts identified in long-read data. GENCODE transcript type are color-coded with red (protein-coding), blue (processed) and gold (Havana-merged). Coding regions are highlighted by filled color in the GENCODE annotation tracks. (**d**) Side-by-side comparison of *IMMT* isoforms expression in *RBM20* R634Q, P633L mutant and WT. Two novel transcript isoforms in red, IMMT-SL1 and IMMT-SL2, were only expressed in WT samples. Known GENCODE transcripts are in green and raw long-reads are in blue. The mis-spliced exon 6 is indicted by *. IMMT-205 and IMMT-208 were omitted due to low read count (1 in all samples).

Notably, not all 107 genes with differentially expressed transcript isoforms had significant changes in total gene expression level, as assessed using standard differential gene expression analysis (Methods, Figure 4b). Among the 107 genes, 27 expressed a single isoform, and of the remaining 80 genes with multiple isoforms, 69 genes displayed total gene expression differences. Among these 69 genes, 60 genes showed differences in major isoform expression (>50% of total transcripts of gene), while 9 genes differentially expressed only minor isoforms. Eleven genes with multiple isoforms showed no change in total gene expression but differentially expressed one or more isoforms. When assessing transcript isoform usage, 8 out of these 11 genes showed significant difference in isoform ratios between mutant and wildtype (Fisher’s exact test, adjusted P-value<0.005, Methods, Supplementary Fig. 20), clearly suggesting complex regulation at the level of splicing, independent of promoter or isoform stability differences since total gene expression was not affected (Supplementary Table 5). This complex isoform regulation demonstrates that measuring differential RNA expression at the gene level and not at the isoform level can fail to detect significant expression changes in specific isoforms, and approaches that do not require prior annotation are advantageous. Two prominent examples of genes with differential expression of specific transcript isoforms (and not total gene expression) are *TPM1*^37^ and *IMMT*^*38*^, both previously reported to be mis-spliced from short-read sequencing of rats and/or human cardiac transcriptomes. Our long-read data showed clearly that not all isoforms are equally affected in the mutants (Figure 4b). For *IMMT*, our percentage of exon inclusion (PSI) analysis on short-read data identified only exon 6 as differentially spliced between wildtype and the mutants, as reported in previous studies^37,38,41,42^ (Figure 4c, Supplementary Text). However, the effect of mis-splicing on full-length transcript isoforms was not captured by previous short-read analyses^37,38^.

Specifically, it remained unclear how many and which transcripts are affected by RBM20 mutations. For *IMMT*, based on the reference GENCODE annotation, it’s logical to hypothesize transcript IMMT-205 as the only affected isoform, as this is the only annotated transcript of *IMMT* without exon 6. Differential expression analysis on transcript quantification based on short-read data indeed reported IMMT-205 as the only significant hit for IMMT (Methods). This is misleading because our long-read based approach could clearly identify two novel *IMMT* transcript isoforms, IMMT-SL1 and IMMT-SL2, that lack exon 6 compared to their corresponding known transcripts, IMMT-207 and IMMT-204. These two isoforms are expressed in the wildtype, absent in R634Q and only marginally expressed in P633L, in contrast to three isoforms that are not differentially expressed (Figure 4a and d). Only one read was identified for IMMT-205 in all samples. Based on domain analysis, we found that the lack of exon 6 may result in proteins with new functions, as the absence of 32 amino acids encoded by this exon disrupts one existing coiled-coil domain and an intrinsically disordered region in the protein (Supplementary Fig. 21). Thus, the discovery of novel IMMT isoforms illustrates the utility of full-length transcriptome identification and quantification without prior annotation.

In conclusion, we present a genome-wide analysis of full-length transcript isoforms of human cells and its differential analysis in the context of splicing mutations in a critical heart gene, *RBM20*. While differential exon usage has been described in splicing-deficient *RBM20* mutations^37,38,40-42^, we discovered complex isoform deregulation on select transcripts, that likely reflects complex combinations of promoter activity, isoform stability and alternative splicing changes on individual genes in RBM20 mutants. Our FulQuant workflow allows direct identification and quantification of transcript isoforms without prior annotation. Exemplified by two novel *IMMT* transcript isoforms that are alternatively spliced, annotation-free full-length isoform sequencing is critical for identifying the existence of many new, informative transcript isoforms that were previously missed. These observations, combined with the overall, intricate complexity of the human transcript isoform landscape (depicted in Figure 1 and in the genome browser), demonstrate the critical need for more widespread adoption of long-read sequencing for transcriptome studies to provide context to previously identified alternatively spliced exons. Knowing the full-length isoform is essential to understanding the functional product of an expressed gene. Our workflow to quantify and compare full-length transcript isoforms directly, including various classes of novel transcripts, can be readily extended to other cell types and tissues, and the many diseases in which aberrant splicing has been proposed to play a role.

## Acknowledgements

We thank Lars Velten, Aaron Brooks, Markus Grosch, Benedikt Rauscher, Julia Kornienko and Sibylle Vonesch for helpful discussion and critical comments on the manuscript. We thank Life Science Editors for their editorial assistance. This study was supported by the National Natural Science Foundation of China (Grant No: 81870187) and National Key R&D Program of China (2017YFC0908405) to W.W., the Chan Zuckerberg Foundation (2019-202666) to L.M.S., the Steinmetz Cardiomyopathy Fund to L.M.S., and the Walter V. and Idun Berry Fellowship to J.W..

## Author contributions

W.W., C.Z., J.W. and L.M.S. conceived the project. J.W. developed the full-length sequencing method and performed the sequencing experiments. H.S. performed the short-read analysis. F.B. generated the iPSC-CMs. C.Z. established the FulQuant method with inputs from W.W.. C.Z. and W.W. performed the analysis on long-read data. B.M. interpreted RBM20 data on clinical perspectives. C.Z., J.W., W.W., and L.M.S. wrote the manuscript.

## Supplementary information

is available for this paper.

## METHODs

### Genome editing in human iPSCs

Methods for genome editing and iPSC-CM generation are detailed in our publication^42^. Briefly, we obtained human iPSCs from the Stanford Cardiovascular Institute Biobank. Annealed (T4 ligation buffer, NEB) and phosphorylated (T4 PNK, NEB) guide RNA (gRNA, designed using tools at http://crispr.mit.edu) oligonucleotides were introduced into the BbsI sites of the pSpCas9(BB)-2A-GFP plasmid and transformed into STBL3 *E. coli* cells and verified via Sanger sequencing. One day after plating the cells into Matrigel coated 6-well plates at low density in Essential 8 (E8) media, media was changed to E8 with Rock inhibitor (Tocris Cat. No. 1254). For each well, CRISPR/Cas9 vector (1μ, pSpCas9(BB)-2A-GFP) and 4μg of single-stranded DNA donor (Supplementary Information) were introduced into the cells via transfection with Lipofectamine 3000. GFP+ cells were isolated 36-48 hours after transfection using a FACSAria IIu (DB Biosciences) flow cytometer with a 100-μm nozzle, maintained in E8 with Rock inhibitor for the first 3 days with an initial density of 2-3×10^3^ cells/well, and subsequently transferred to regular E8 until the colonies size reached ∼0.5 mm. Individual iPSC clones were isolated and re-plated in a well of a 24-well plate in E8 with Rock inhibitor. The homozygous edits were confirmed by sequencing after PCR amplification of the target genomic region (Supplementary Information). The R634Q cells will be made available upon request.

### iPSC-CM generation

iPSCs were differentiated into cardiomyocytes as a monolayer through the modulation of WNT signaling. iPSCs were plated at low density on Matrigel coated plates to have them 70-80% confluent after 4 days (Day 1 of differentiation). Differentiation was induced with 3ml of RPMI 1640 (Life Technologies 11875-093) with 1X B27® Minus insulin (Life Technologies 0050129SA), supplemented with 6μM CHIR (TOCRIS 4953). On day 6, media was replaced with 3ml of RPMI 1640 with 1X B27® Minus Insulin, supplemented with 5μM IWR. From day 8, cells were kept in RPMI 1640 (Life Technologies 11875-093) with 1X B27® Serum-Free Supplement (Life Technologies 17504-044). From day 12 to day 15 cells were treated with RPMI 1640 no Glucose (Life Technologies 11879-020) with 1X B27, then allowed to recover for 2 days in RPMI 1640 with 1X B27, and subsequently replanted at a density of 3Mi cells/well. Cells were then maintained in RPMI 1640 with 1X B27 for 4 more weeks before being collected for the experiment.

### Reverse transcription

RNAs were cleaned and treated with DNase using ZymoResearch RNA Clean and Concentrator-5. A total of 4.5 μL mixture containing 5 ng of purified iPSC-CM RNAs with 2.8% ERCC and 3% Sequins Version A, 1μL of 10μM oligodT (/5SpC3/A*A*G*CAGTGGTATCAACGCAGAGTACTTTTTTTTTTTTTTTTTTTTTTTTTT TTTTVN), 1μL of 10mM dNTP, 0.1μL of Takara recombinant ribonuclease inhibitor (40U/μL) were incubated at 72 °C for 3min, 4°C for 10min and 25°C for 1min. Another 5.5μL mixture containing 2 μL of 5x SuperScript II buffer, 2μL of 5M betaine, 0.5 μL of 100 mM DTT, 0.5 μL of SuperScript II (200 U/μL), 0.25 μL of Takara recombinant ribonuclease inhibitor, 0.1 μL of 100 μM TSO, 0.09μL of nuclease-free water, 0.06 μL of 1M MgCl_2_ were added. The total 10 μL RT reaction were incubated at 42°C for 90 min, 10 cycles of 50°C for 2min and 42°C for 2 min^4^. The reaction was then stopped by incubation at 70°C for 15 min and held at 4°C. The reaction was then treated with 1μL of 1:10 dilution of NEB RecJf (Catalog # M0264S) at 37°C for 30min and 65°C for 20min.

### PCR amplification of full-length cDNA

25μL of 2x NEB Q5 HotStart master mix, 0.25 μL of 10μM PCR primers (ONT_Index1_ISPCR and ONT_Index2_ISPCR)^5^ and 14.5 μL of nuclease-free water were added to the RT reaction. The PCR were performed as follows: 98°C for 30s, 20 cycles of 98°C for 10s, 67°C for 15s, and 72°C for 6min, then 72°C for 2min, and hold at 4°C. PCR reactions were purified using 0.75 volume of Ampure XP beads and eluted with 25 μL of nuclease-free water. Full-length cDNA were analyzed on Agilent Bioanalyzer using High Sensitivity DNA chips.

### Oxford Nanopore Sequencing

1μg of cDNA were ligated with ONT sequencing adaptor as described by the manufacturer with modifications in the following steps: 1) End repair and dA tailing; 2) Ligation of ONT sequencing adaptor; 3) Addition of sequencing tether. Sequencing was performed using either R9.4 and R9.5 flow cells on the MinION device. Data were recorded with ONT’s MinKNOW software for 48h. Basecalling was performed using ONT’s Albacore software (version 2.1.3) with the options ‘--disable_filtering --kit SQK-LSK108’ and corresponding flow cell version (Supplementary Table 1).

### FulQuant for full-length transcript isoform quantification

We assessed the dependency of percentage of alignment identify (PID) on mean read quality (mean_qscore defined by ONT) and determined the mean read quality score cutoff 6 where over 80% reads have a PID > 80% (Supplementary Fig. 2). After removing low quality reads, we trimmed sequencing adapters ISPCR (AAGCAGTGGTATCAACGCAGAGTAC) and polyA sequences (>=10 consecutive A/T with one mismatched allowed) at both ends (200 bp window) of the reads. Reads having adapter sequence at one end and polyA sequence at the other end were considered full-length. We aligned the reads against the human genome (GRCh38) using minimap2 (version: 2.9-r751-dirty) with the parameters: ‘-K500m --secondary=no -a -x splice --splice-flank=yes’. Alignments meeting the following criteria were removed from downstream analysis: i) consist of supplementary alignments, ii) size of soft clipping at either ends greater than 30 bp, iii) insertion size near splice junctions (identified by cigar N) greater than 5 bp, iv) unspliced alignment (singleton).

The transcriptome annotation was performed on data for WT and R634Q. We identified tag splice sites for intron-exon and exon-intron junctions using a clustering method previously described^45^ with the parameters ‘HFWINSIZE=5 DISTHRES=8 PTHRESHOLD=3 SUMHOLD=5’. Reads sharing the same tag splice sites were collapsed into consensus transcript isoforms. We include the terminal exons but do not consider transcript variation at 5’ and 3’ ends due to high error rate at both ends of the read. In fact, for the terminal exons, we annotated their splicing sites precisely, but not their transcript start (TSS) and stop (TES) sites. We used the median 5’ and 3’ positions of all reads in one transcript isoform as its TSS and TES sites. The following rules were used to remove false positive transcripts: 1) read count >3, 2) >40% full-length reads, 3) transcript termination site greater than 10 bp from known genomic polyA sequence, 4) strand information is available. Furthermore, we required in each gene locus that all transcripts: 1) had more than 3% read count of the most abundant transcript, 2) were not results of misalignment, 3) were not 5’ truncation products of any identified transcripts with higher counts, unless the count is higher than 25% of the longer transcript. The code used to perform the above steps are attached in a supplementary file with detailed annotation. For the final annotation, we also removed transcripts that either had only 2 exons or may be a truncation product of a known transcript.

We compared our annotation with GENCODE human comprehensive annotation (V24) to classify transcripts as known and novel. Filtered transcripts mapping to known GENCODE annotation were rescued. We further annotated the novel transcripts into novel exons, novel combination, exon skipping, intron retention, alternative splice sites, and multiple events. A detailed definition of each novel transcript class can be found in the supplementary information and code.

### Exon connectivity analysis

We first defined exons that were present in all transcripts as constitutive. For all exons pairs within a gene, we calculated the number of reads for when: 1) both exons are present, 2) only exon 1 is present, 3) only exon 2 is present, 4) both exons are not present. Using this count matrix, we tested their co-association using fisher’s exact test following p-values adjustment with BH method. Distal pairs were defined as exon pairs that were at least one constitutive exon apart.

### Predicating nonsense-mediated decay events

For all novel transcripts that include a known start codon defined in the GENCODE annotation, we calculated the distance between the first stop codon and the last splice junction on the transcript. We defined transcripts as NMD products if this distance was greater than 55 bp^46^.

### Short-read data analysis

After mapping reads to GRCh38 using STAR v2.5.1b, gene expression levels were determined with featureCounts v1.6.0. For each exon, we counted inclusive and exclusive reads as reads that include the exon of interest and reads that include both the upstream and the downstream exon but not the exon of interest, respectively. Percentage of spliced-In (PSI) representing the ratio between inclusive reads over the sum of inclusive and exclusive reads was calculated. Transcript quantification was performed using Kallisto (0.46.2) with either GENCODE or our annotation as input annotation for index.

### Gene and transcript isoform level differential expression analysis

Transcript quantification was achieved by counting number of reads sharing the same splice sites of that transcript. Gene quantification was achieved by summing transcripts counts in that gene locus. We compared CRISPR generated R643Q and P633L mutant samples to samples without CRISPR editing treatment (WT_NC) and non-edited samples with CRISPR editing treatment (WT), whereby standard differential expression analysis was performed using DESeq2 for both genes and transcript isoforms. Due to the limited sequencing depth, we only considered genes and transcripts isoforms with more than 10 read counts in total in all samples for the test for the long-read data. Significant candidates were defined using a BH-adjusted p-value cutoff of 0.001 for the comparison between mutants and WT. For short-read data, same testing procedure with DESeq2 was applied to transcript and gene quantification obtained using Kallisto without any filtering. GO enrichment analysis was performed using PANTHER via the web interface at http://geneontology.org with the expressed genes as the reference list to control for background. To assess difference in transcript usage between mutant and wildtype, we summarized biological replicates and performed Fisher’s exact test for the 80 candidate genes with one or more differentially expressed isoforms. P-values were adjusted using BH procedure.

### Validation of novel splicing events using fragment analysis

We used fragment analysis to validate 33 novel splicing events (Supplementary table 6 and Fig. 10). Oligo primers targeting the novel splicing events were designed such that amplicons with and without the splicing events would display detectable size differences on the gel. All primer sequences can be found in Supplementary table 6. We performed standard RT-PCR and analyzed the fragment sizes using standard Bioanalyzer profiling. Only profiles contained band(s) with intended size(s) were considered as positive validation. For 7 unsuccessful cases, 3 cases didn’t yield any PCR product and 4 cases contained fragments, which were at least 20 bp off the predicted sizes.

### Data availability

The long-read data reported in this study have been deposited in ArrayExpress under accession number E-MTAB-7334.

The short-read data reported in this study have been deposited in Sequence Read Archive (SRA) under accession number PRJNA579336^42^.

Our genome-wide full-length isoform dataset is available as a custom genome browser at http://steinmetzlab.embl.de/iBrowser/.

### Code availability

R code to perform the essential steps of transcripts filtering is currently available on our website http://steinmetzlab.embl.de/Http_sharing/czhu/nanopore_tx_filtering.zip and will be made available on GitHub https://github.com/czhu/FulQuant upon publication.

