## Supplementary Information for "Single-molecule, full-length transcript isoform sequencing reveals disease mutation-associated RNA isoforms in cardiomyocytes"

### Supplementary Text

#### Detailed analysis of previously unidentified transcripts

Based on the identified novel events, we categorized 5,740 novel transcript isoforms into following categories: transcripts with novel exons (NE), alternative splice sites (ASS), novel intron retention (IR), novel exon skipping (ES, 12.7%) and unannotated combinations of known exons (NC). NE are defined as exons with a novel combination of the splice sites that was unannotated in GENCODE annotation. It can be either a new combination of known splice sites (as "+" cases in Figure 1) or previously unidentified splice sites (as "#" cases in Figure 1). We found 288 of the novel exons that are surrounded by novel splice sites, 99.5% of which matched canonical splice-acceptor or donor motifs (GT/AG), indicating that they are *bona fide* splice sites. ASS events are defined as exons with one splice site known, and the other unknown. It can be either an extended or shortened version of a known exon. A number of transcript isoforms (140 or 2.4%) contain more than one novel event present simultaneously, indicating various complex splicing patterns. Potential read through transcripts were labeled as dubious due to possible alignment issues of repetitive sequences.

While 116 novel transcripts are not in any known gene loci, thus likely to be non-coding, the remaining of the novel transcripts could potentially encode proteins with novel functions. To investigate their coding potential, we first investigated if these novel transcript would be subjected to non-sense mediated decay (NMD, Figure 2f). Over 60% of novel transcripts with NC and ES events are non-NMD products. Novel transcripts with alternative splice sites (ASS) have a 42% non-NMD product proportion. As expected, novel transcripts with IR have the least non-NMD proportion (16%), suggesting that the majority of them constitute immature, unprocessed mRNAs. Furthermore, 119 novel transcripts are products from known protein-coding transcripts with one additional novel exon spliced in. 26 of these 119 novel transcripts are likely to encode new peptide sequences.

#### Comparison to CHESS transcriptome annotation

When comparing our annotation with the CHESS (v2.2) transcriptome annotation, we found 29,833 of our transcripts are also annotated in the CHESS database (Supp. Fig. 8). The highest agreement is found for the GENCODE annotated transcripts (REF). In total, 47.2% (2712 out of 5740) of novel isoforms we identified in our study when comparing to GENCODE are also present in CHESS annotation. Notably, over 50% of our new transcripts with novel combination of existing exons are independently validated by the CHESS database. We found the lowest agreement for the intron retention transcripts. We believe this is possibly due to the limit of assembling long fragments using short-read RNA-seq, as the CHESS annotation is based on short-read RNA-seq experiments

#### Comparison to FLAIR identification and quantification results

To benchmark our FulQuant pipeline against existing software for transcript identification and quantification, we analyzed our long-read dataset using FLAIR (https://github.com/BrooksLabUCSC/flair, v1.4.0) using the default settings. FLAIR reported 281 transcripts for the synthetic control SEQUIN dataset, only 87 of these were actually present in the SEQUIN annotation, thus resulting in a false positive identification rate of 70% (1-87/281). As comparison, our FulQaunt pipeline has an overall false positive rate of 3%, illustrating the importance of filtering out false positive transcripts.

When running on our human dataset, FLAIR reported 90822 multi-exonic transcripts, only 27545 of which agree with GENCODE annotation. Among these known transcripts, 25682 are also reported in this study. However, the overlap for the novel transcripts are much smaller between FLAIR and FulQuant. Only 3073 transcripts reported by FLAIR agree with our novel transcripts and other 60204 novel transcripts are likely to be false positives based on FLAIR's performance with synthetic control SEQUIN.

#### Comparison to PSI-analysis results on short-read data

PSI-based analysis on the short-read data identified 22 candidate genes including IMMT, SYNPO and TPM2, which we also identified in our long-read analysis. However such PSI analysis doesn't pinpoint to the exact transcript isoforms with differential expression. We found 104 and 19 candidate genes are specific to the long- and short-read data, respectively. Comparison with hits from previous studies (Guo et al., Nature Medicine, 2012; Maatz et al., The Journal of Clinical Investigation, 2014) yielded similar overlaps (Supplementary Figure 20).

#### Comparison to Kallisto quantification results

We performed differential transcript expression analysis using Kallisto quantification. Using GENCODE annotation, such analysis doesn't identify novel transcripts, such as two novel IMMT transcripts we highlighted. Using our annotation as input for Kallisto, we can confirm differential transcript isoforms identified by our long-read dataset

(Supplementary Figure 15 and 16). The major difference likely resulted from the significant differences in sequencing depth between the two datasets, with long-read data having about 1-2 million effective reads per sample after stringent filtering compared to ~ 200 million pair-end reads per sample in the short-read data.

#### GO enrichment analysis

The 11 genes we identified with specific isoform deregulation in the absence of overall expression changes in the RMB20 mutant versus WT cells belong to genes enriched in the Gene Ontology (GO) category of muscle thin filament tropomyosin component (adjusted $P<0.005$), suggesting that they may have a disease-relevant function.

#### Supplementary Figures and Legends

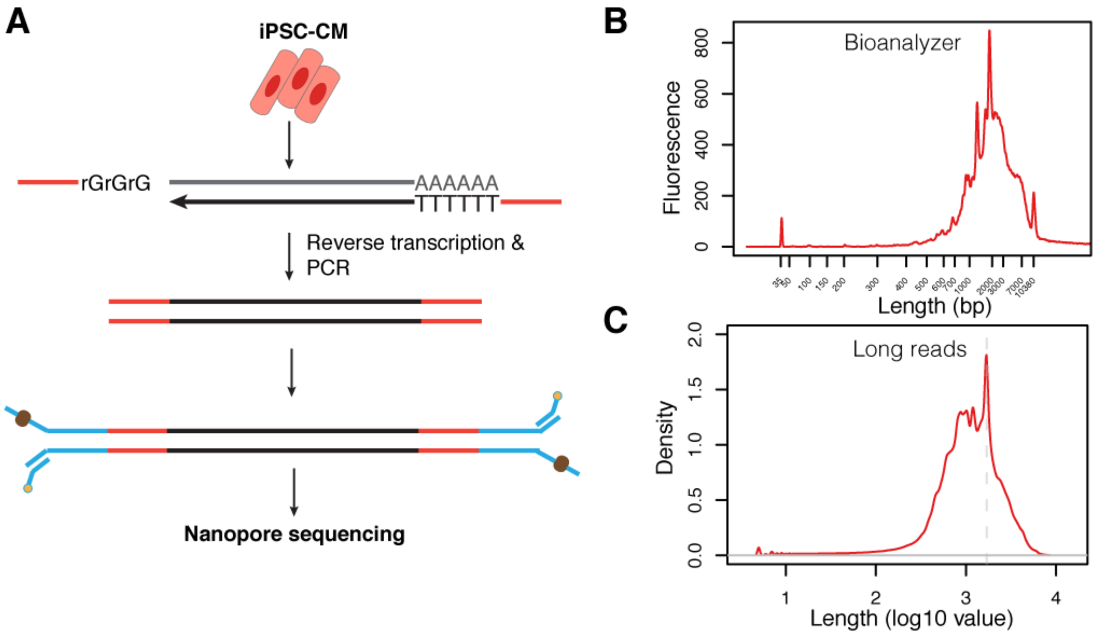


**Supplementary Figure 1. Workflow of full-length cDNA sequencing and analysis protocol.** (a) Experimental pipeline for generating full-length cDNA molecules. (b) cDNA length distribution by Aglient High-sensitivity Bioanalyzer. (c) ONT reads length distribution of the same sample.

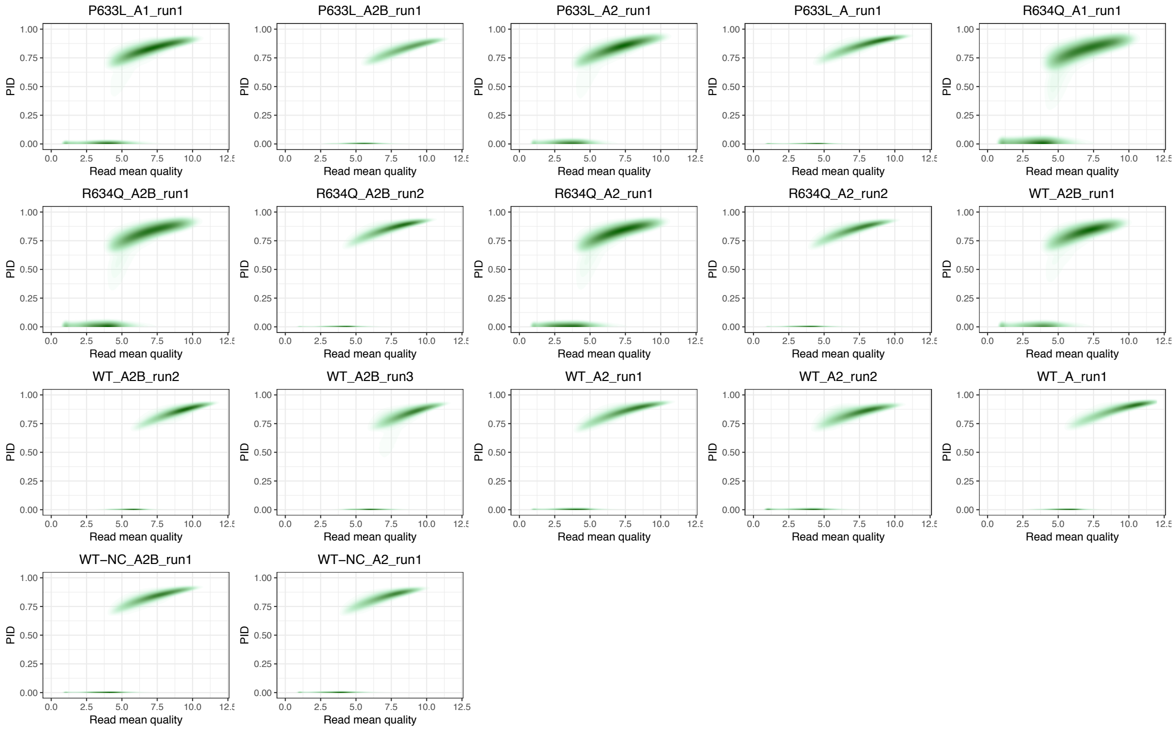


**Supplementary Figure 2. Relationship between percentage identity (PID) versus mean read quality for each Nanopore sequencing run.** For each read, percentage identity (percentage of aligned read over total read length) was plotted against mean read quality as reported by basecaller Albacore. This plot was used to determine the read quality cutoff of 6, with which ~ 80% PID was achieved on average.

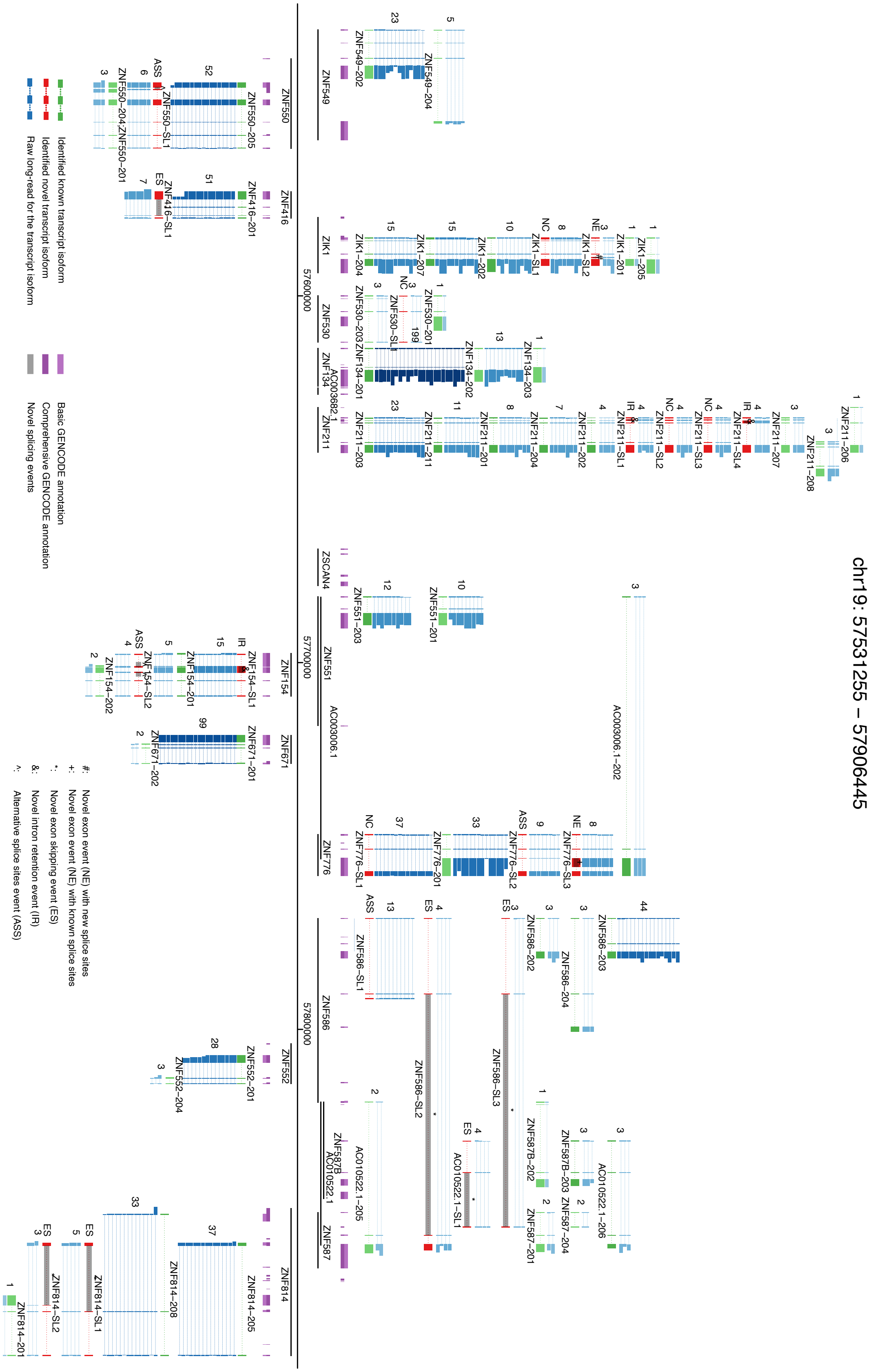


**Supplementary Figure 3. Complex landscape of full-length human iPSC-CM transcriptome**. This figure shows the same region of chromosome 19 (chr19:57531255-57906445) as Figure 1 with representative raw reads for each isoforms rendered. Gene loci are presented as a horizontal black line with annotated genes on + or – strands above or below the genome axis, respectively. Collapsed GENCODE comprehensive and basic annotation are presented in dark purple and purple track. Known transcript isoforms are shown as green tracks, and previously unidentified transcript isoforms as red. Known transcript isoforms didn't pass transcript filtering criteria are presented in light green. Raw long-reads for each isoform are shown in blue (darkness correlates with expression level) with numbers of track lines in logarithmic scale of raw read number (left of the tracks). Based on the novel splicing events (location indicated with grey box and text symbols), novel transcript isoforms are categorized as novel exon combination (NC), novel exon (NE), novel intron retention (IR), novel exon skipping (ES), or novel alternative splice sites (ASS).

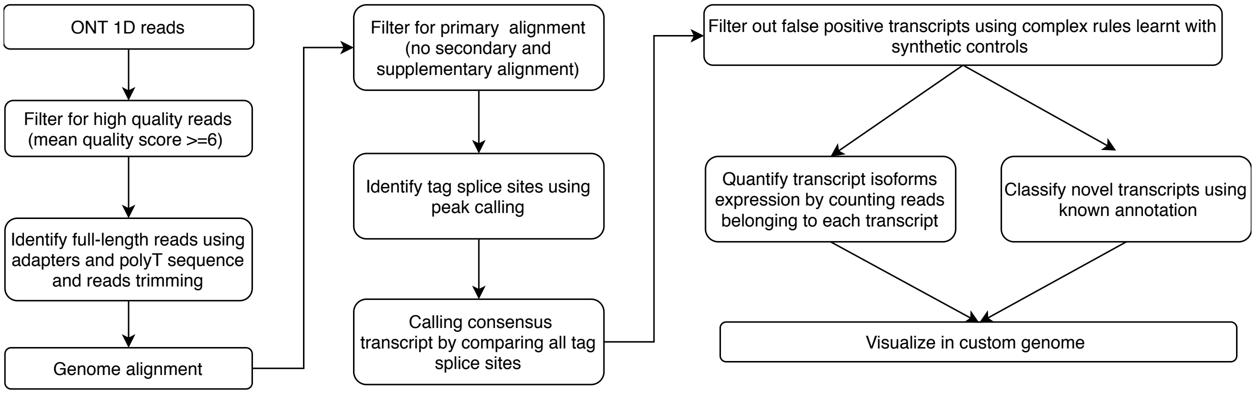


**Supplementary Figure 4. Workflow of the FulQuant computational pipeline for quantifying and visualizing full-length transcript isoforms with ONT 1D reads.** ONT reads were filtered by quality score and sequencing adapters were trimmed. Following alignment to the human genome, reads were grouped based on the sharing of all splice sites. Filtering was performed at both the read and transcript levels. By comparing to the GENCODE annotation we classified transcripts into different groups and subsequently visualize them in a custom genome browser tailored for visualizing long-read data. The FulQuant method is implemented using in-house scripts.

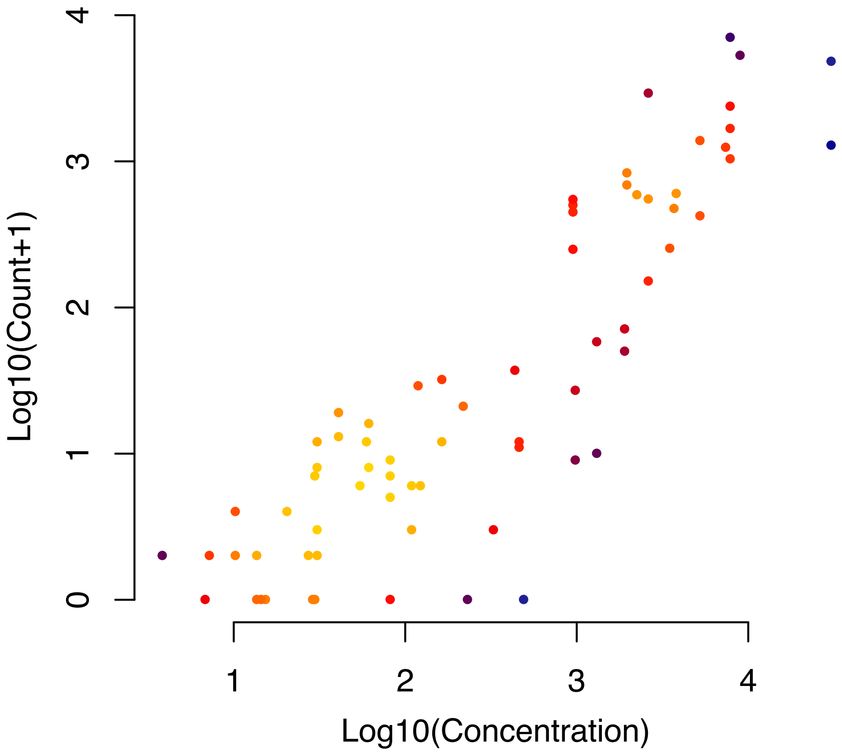


**Supplementary Figure 5. High correlation of sequin read counts to input concentration.** Shown here are data for a sample with a Spearman correlation coefficient of 0.86. All samples share similar correlation (0.8 - 0.9). The color of the points corresponds to data density in the plot.

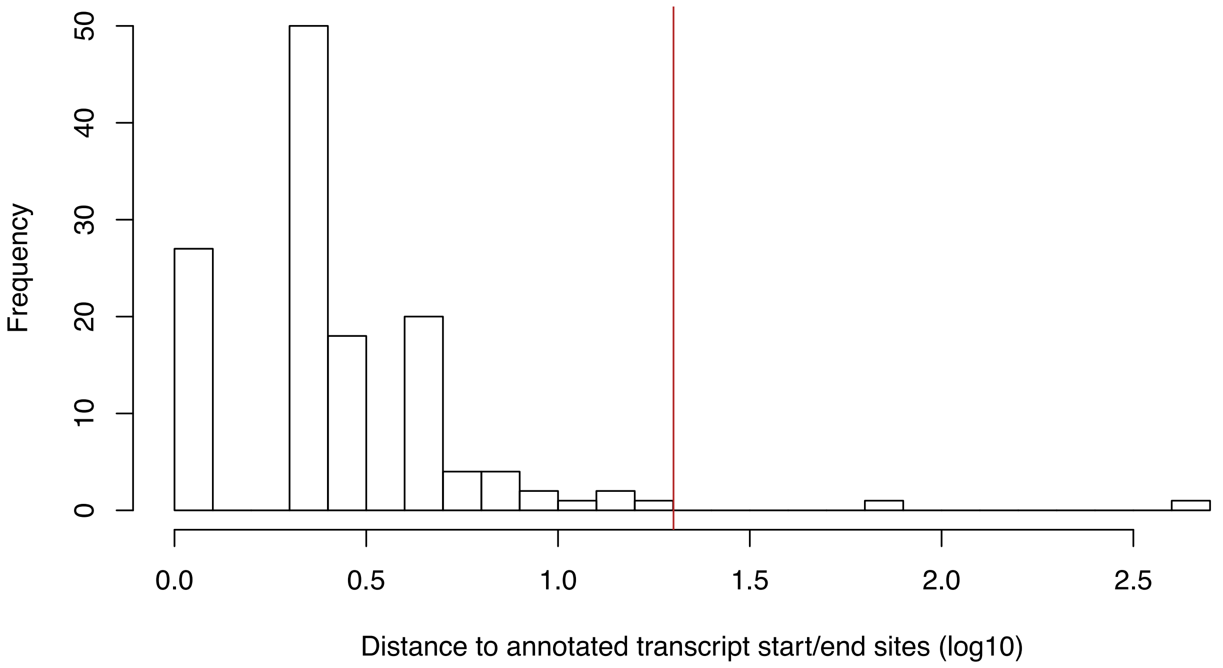


**Supplementary Figure 6. Distribution of distance between identified and nearest annotated transcript boundaries.** Red line denotes 20 bp**.**

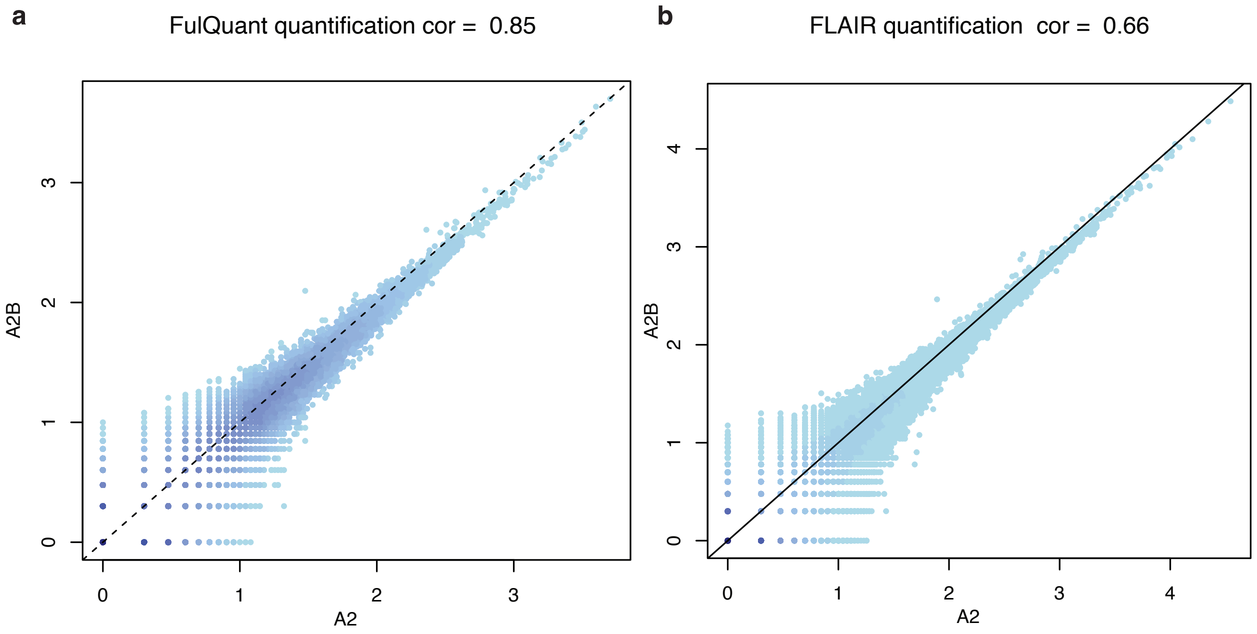


**Supplementary Figure 7**. **Transcript isoform quantification (read count in log10) for biological replicates with a) FulQuant and b) FLAIR software.** Both software were run on the same long-read data generated in this study. FulQuant quantified ~36k transcripts while FLAIR quantified over 90k transcripts, many of which are likely false positives. Shown here are quantified transcripts for two biological replicates of the wildtype sample. Both correlation coefficients (cor) given in the title are Spearman's rank correlation. Color gradient represents point density.

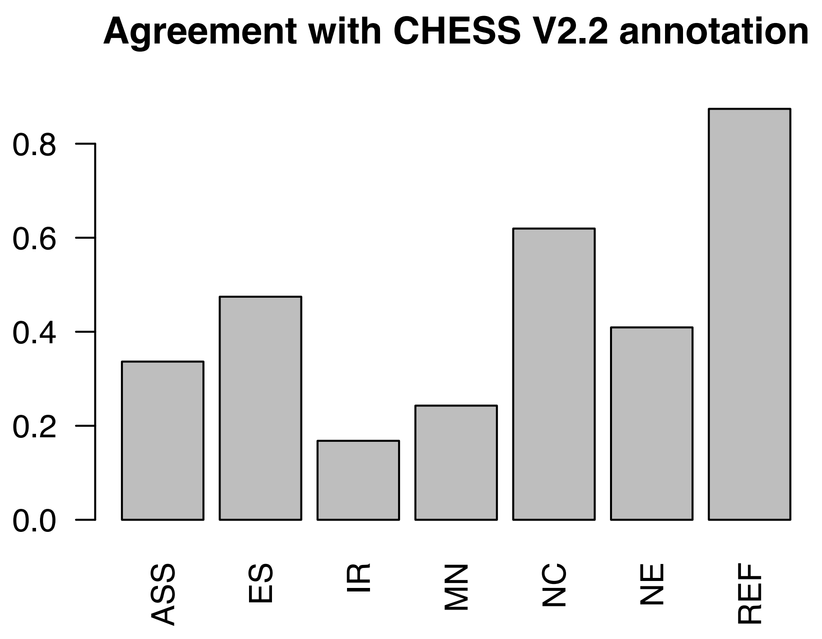


**Supplementary Figure 8**. **Agreement of transcripts identified in this study with the CHESS transcriptome annotation.** Transcripts are considered the same if they share the same set of splice sites. Different types of transcripts include: GENCODE annotated transcripts (REF), novel exon combination (NC) , novel exon (NE), novel intron retention (IR), novel exon skipping (ES), or novel alternative splice sites (ASS). MN denotes multiple novel events in the transcripts.

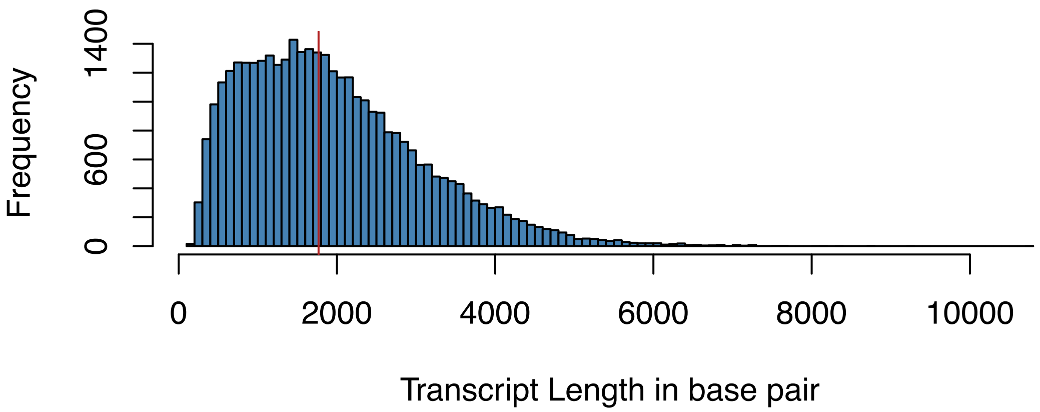


**Supplementary Figure 9. Length distribution of identified transcripts in our long-read data.** Red line denotes the median value 1,768 bp.

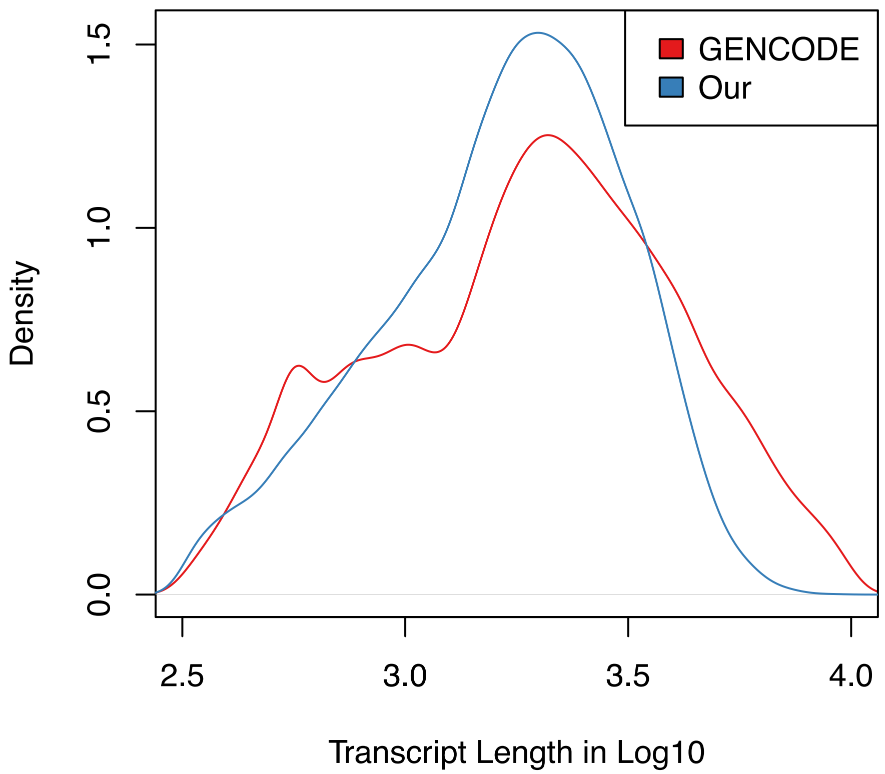


**Supplementary Figure 10. Length distributions of identified transcripts in our long-read data and full-length GENCODE transcripts for protein-coding and lincRNA.**

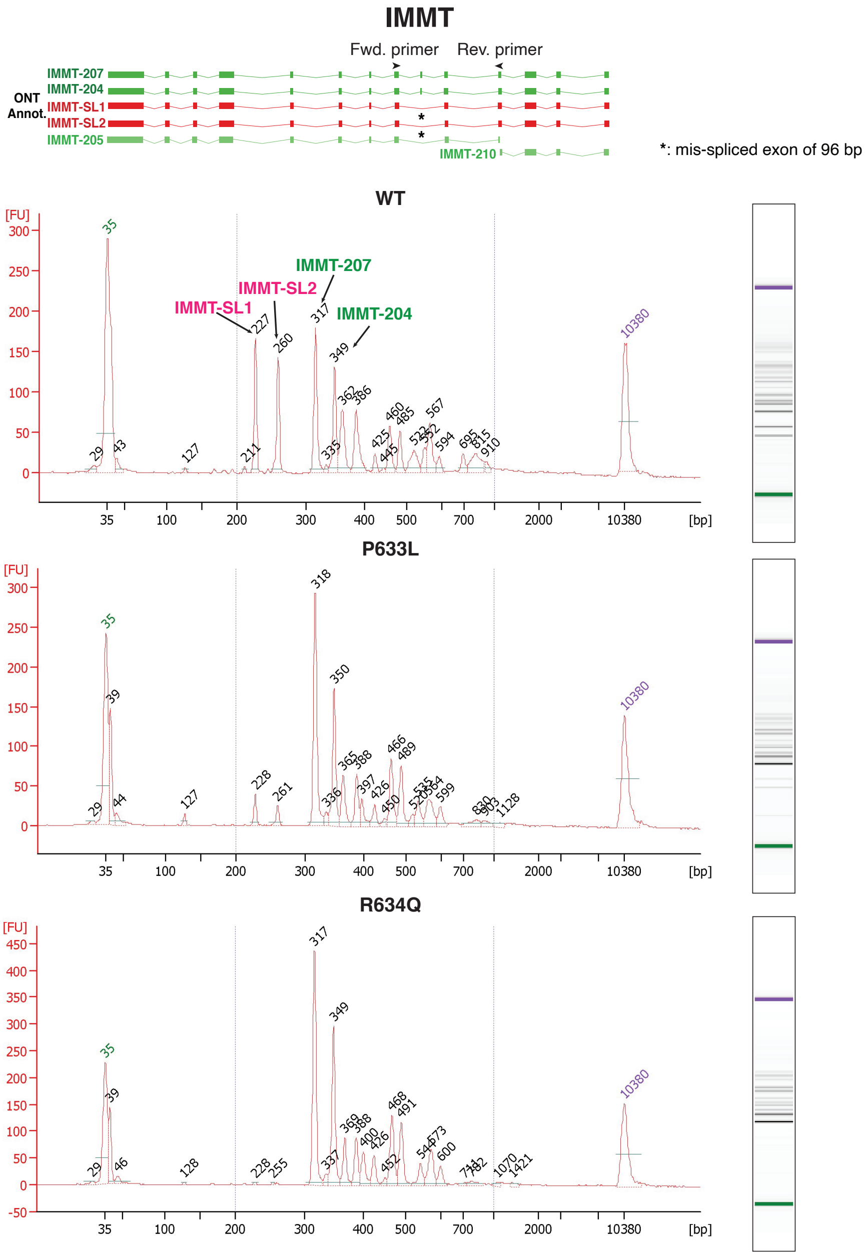


**Supplementary Figure 11**. **Validation of IMMT mis-splicing using fragment analysis.** Primer pair was designed to include the mis-spliced exon 6 (upper panel). Size differences of amplicons with and without the target exon were visualized using Bioanalyazer (lower panel). As expected, novel transcripts IMMT-SL1 and IMMT-SL1 without the 96 bp exon 6 were mostly expressed in WT. Their expression was much lower in P633L and not detectable in R634Q mutant.

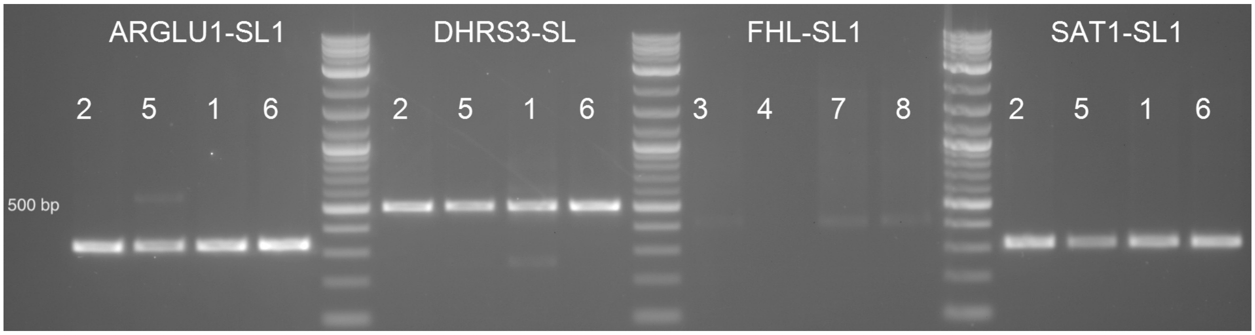


**Supplementary Figure 12**. **PCR validation for intron-retention events.** Intron specific primers were designed using Primer3 with target sizes ranging from 316 to 505 bp. PCR was performed for 4 different templates (Numbers denoting different samples) for each primer pair.

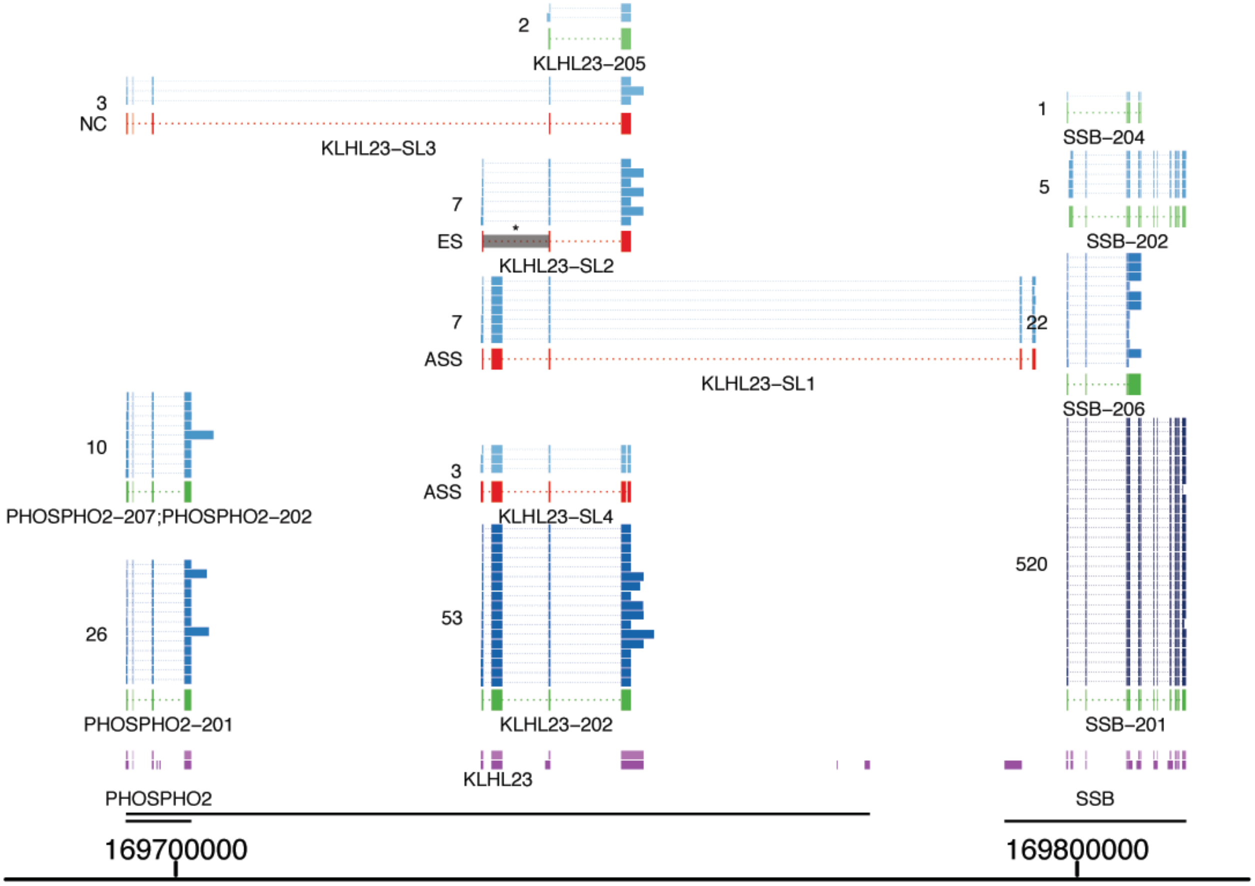


**Supplementary Figure 13. Transcript KLH23-SL1 is a fusion transcript between gene KLHL23 and SSB.**

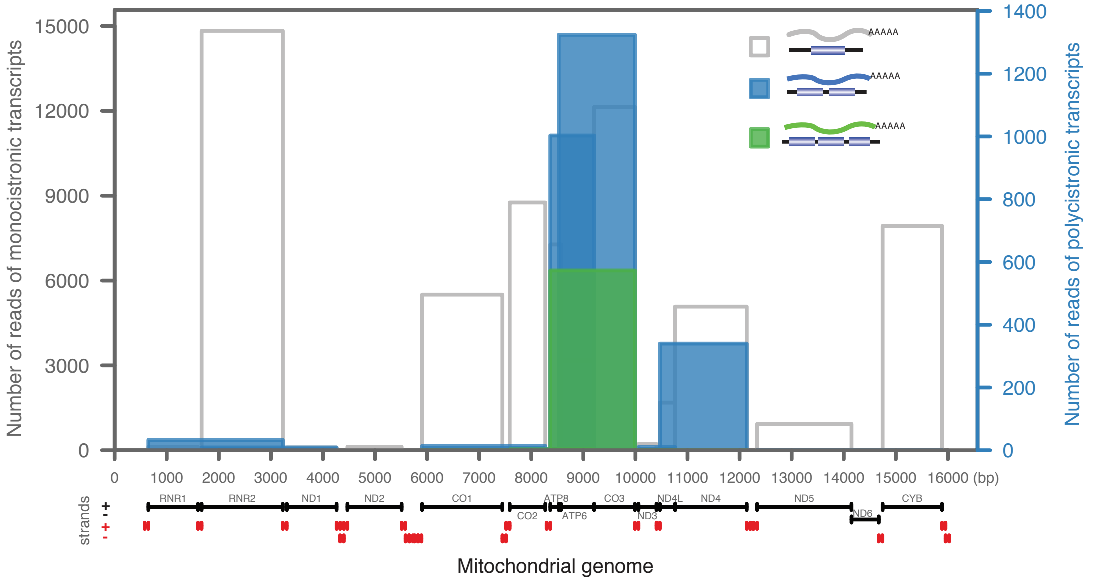


**Supplementary Figure 14. The human mitochondrial genome transcribes large numbers of polycistronic transcripts.** Among all mitochondrial unspliced reads covering the gene coding regions, ~ 6% are polycistronic transcripts. 99% of these polycistronic transcripts are either bicistronic or tricistronic transcripts from regions adjacent to the APT8, ATP6 and COX3 genes, as well as the adjacent ND4L and ND4 genes. Since no tRNA gene is located inside these two genomic regions, this suggests that most mitochondrial transcripts identified in this dataset are processed via tRNA end cleavage.

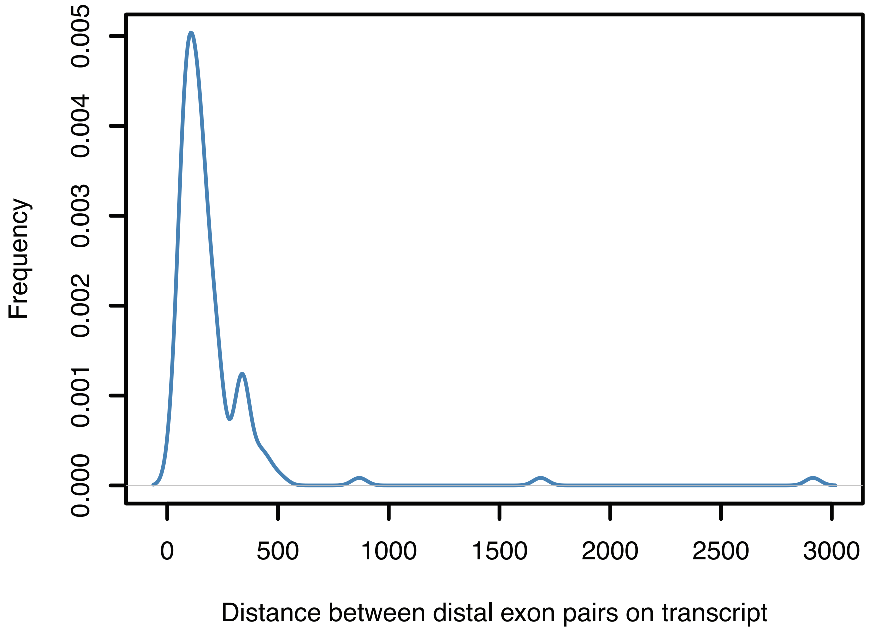


**Supplementary Figure 15. Distribution of distance between distal associated exon pairs.** Most distal pairs are separated by one or two exons.

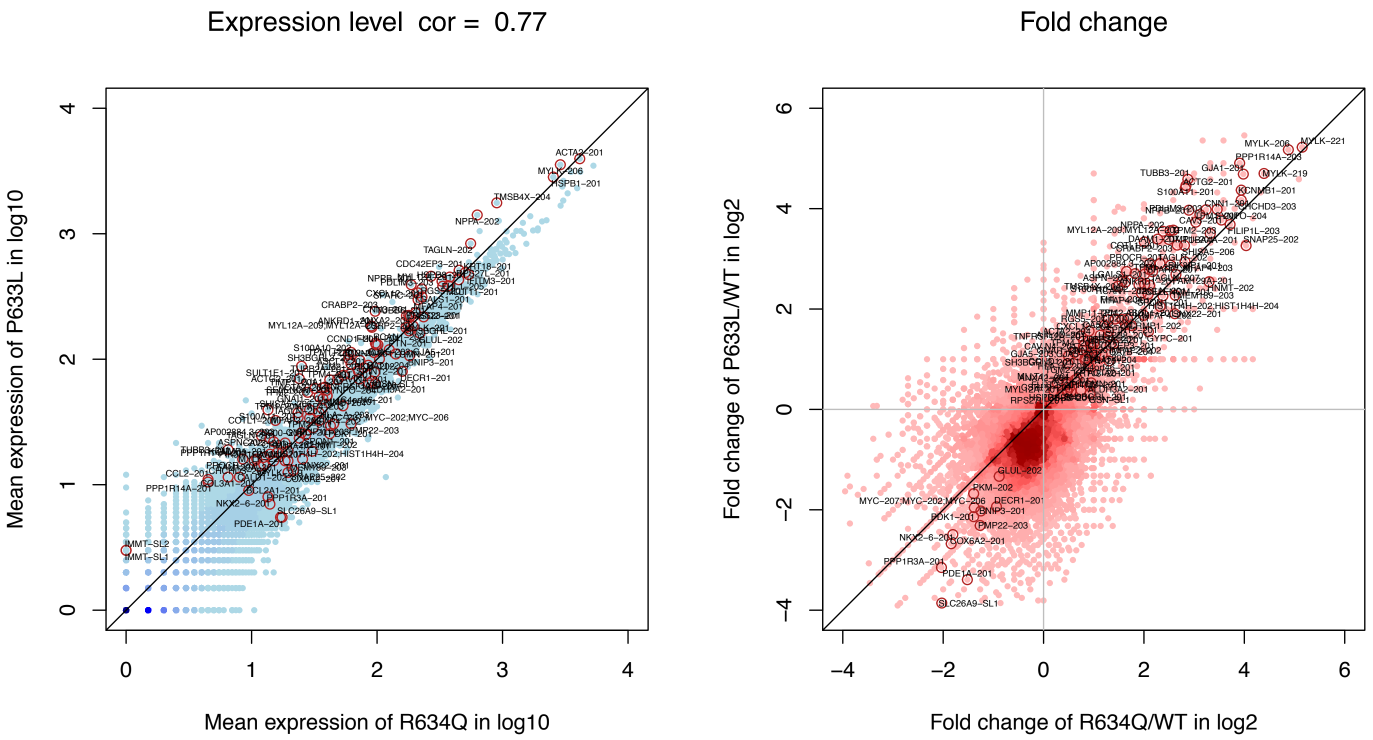


**Supplementary Figure 16. Comparing R634Q and P633L mutants using long-read sequencing.** Transcript isoforms were quantified using FulQuant. Both mutants agree well on average expression levels (left panel, Spearman's rank correlation coefficient 0.77). When compared to wildtype, their effect size show similar trend (right panel). Highlighted points are significantly differentially expressed transcripts reported in our study. Color gradient represents point density.

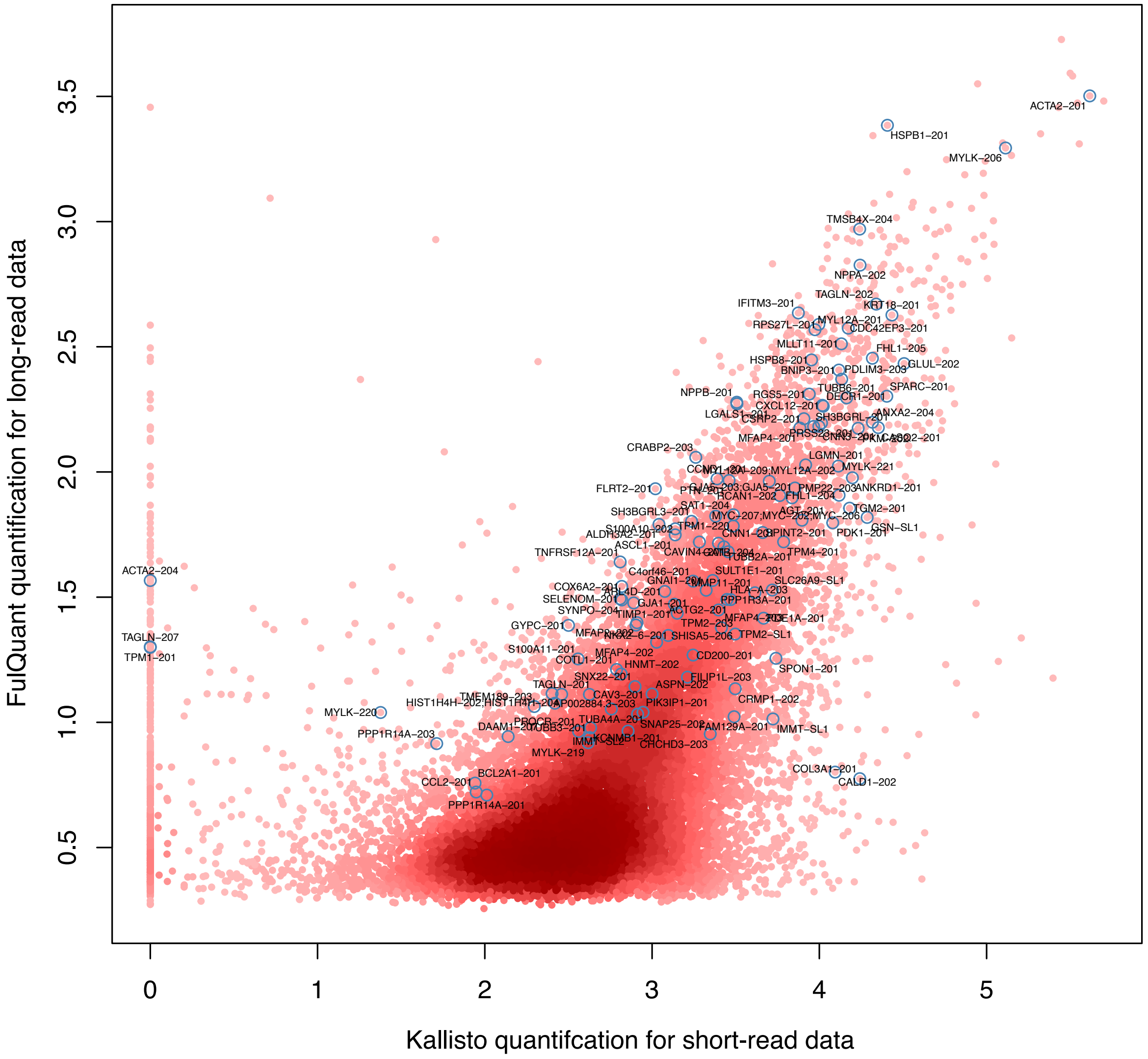


**Supplementary Figure 17**. **Comparison between transcript quantification using long-read and short-read data.** Shown are mean quantification (read count in log2) of all transcript isoforms across all samples for Kallisto (x-axis) and FulQuant quantification (y-axis) . FulQuant and Kallisto were used to quantify transcripts in the long-read and short-read data, respectively. For Kallisto, transcript annotation by FulQuant was used as reference. Despite the differences in sequencing technologies and depth, both datasets agree well (Spearman's rank correlation coefficient 0.63). Highlighted are significantly differentially expressed transcripts reported in our study. Notice Kallisto was unable to quantify several transcripts such as TPM1-201. Color gradient represents point density.

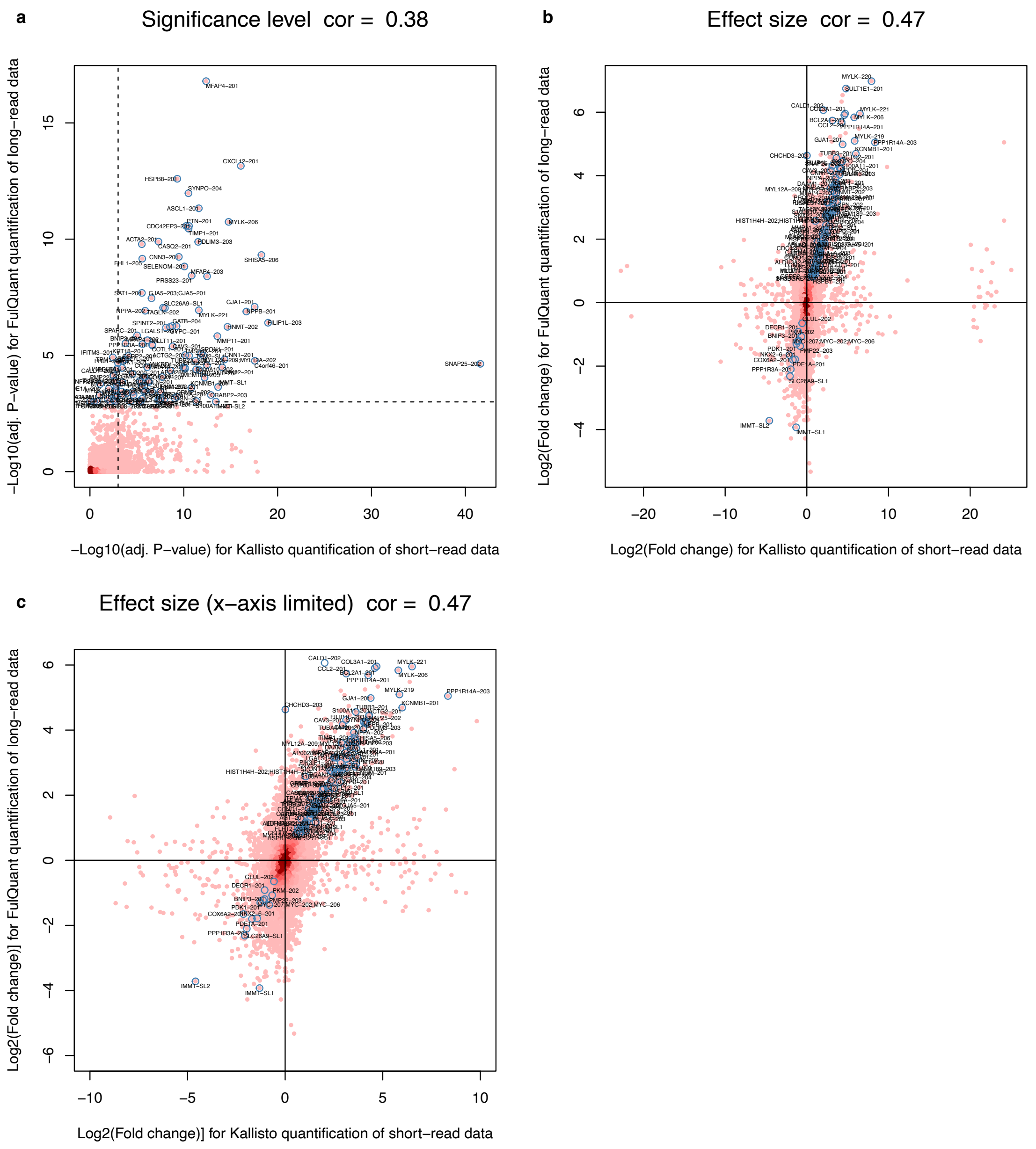


**Supplementary Figure 18**. **Differentially transcript analysis for long-read and short-read quantification.** FulQuant and Kallisto were used to quantify transcripts in the long-read and short-read data, respectively. For Kallisto, transcript annotation by FulQuant was used as reference. Same differential expression analysis was performed on both datasets using DESeq2. The results are visualized for significance level (**a**), fold change (**b**) and fold change with limited x-axis (**c**). Highlighted in all plots are differentially expressed transcripts reported in our study. Both analyses based on long-read and short-read data agree well on the subset of significantly differentially expressed transcripts. Analysis based short-read data had higher sensitivity in detecting expression difference as shown in Figure **a** (dashed line indicates adjusted P-value cutoff of 0.001) due to its much higher sequencing depth. However, some of its effect size estimates are erroneous as demonstrated by extreme points (points around -20 and 20 on x-axis) in Figure **b.** Overall, the fold changes between mutant and wildtype were consistent in both analyses. Color gradient represents point density.

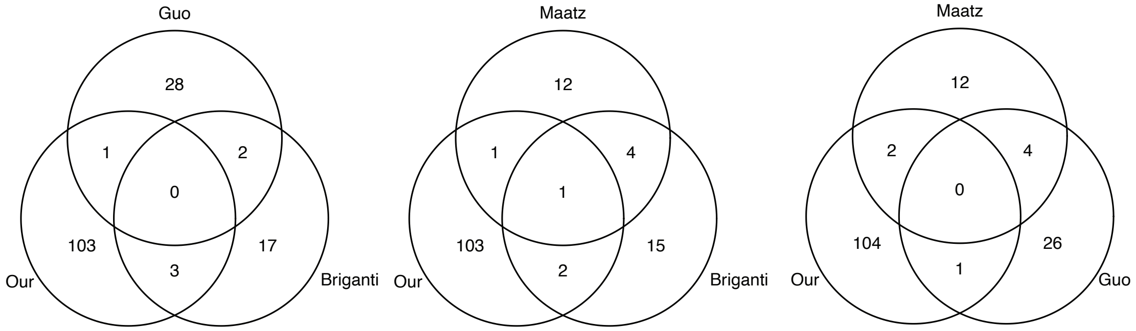


**Supplementary Figure 19. Comparison of RBM20 targets from various studies.** All studies share similar gene hit overlap of 1-4 genes. Our (this study) hits are genes hits from long-read data. Briganti hits are gene hits from PSI analysis of the short-read data in this study. Guo hits are candidate genes reported in Guo et al., Nature Medicine, 2012 study. Maatz hits are candidate genes reported in Maatz et al., The Journal of Clinical Investigation, 2014 study.

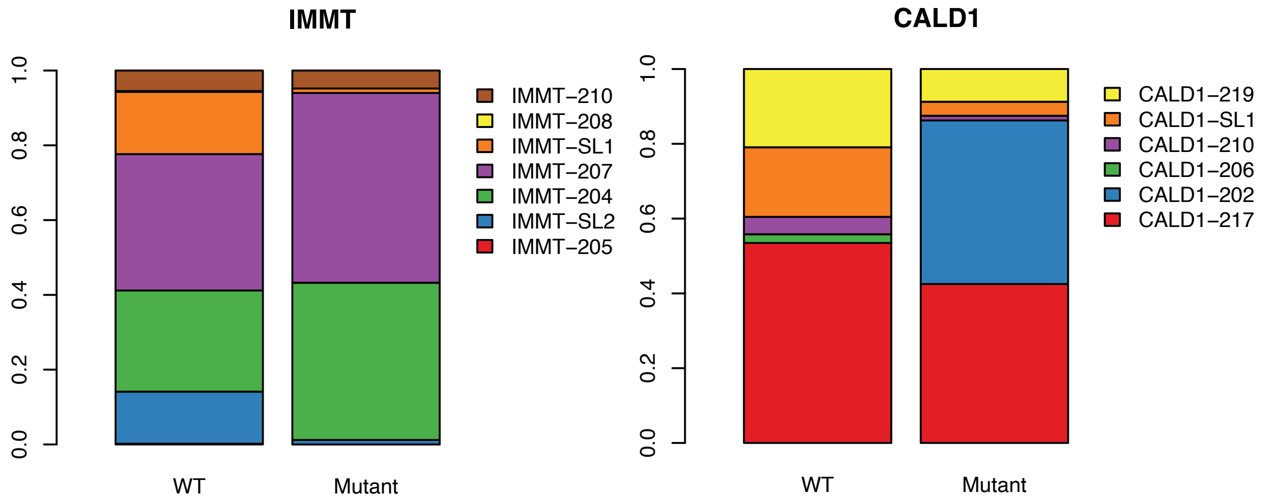


**Supplementary Figure 20. Example genes with significant changes in transcript usage between wildtype and mutant.** Counts for biological replicates were summarized. The ratio of isoform count over the total gene count was plotted. Fisher's exact test was used to assess change in transcript usage for the 80 candidate genes with one or more differentially expressed isoforms. P-values were adjusted using BH procedure.

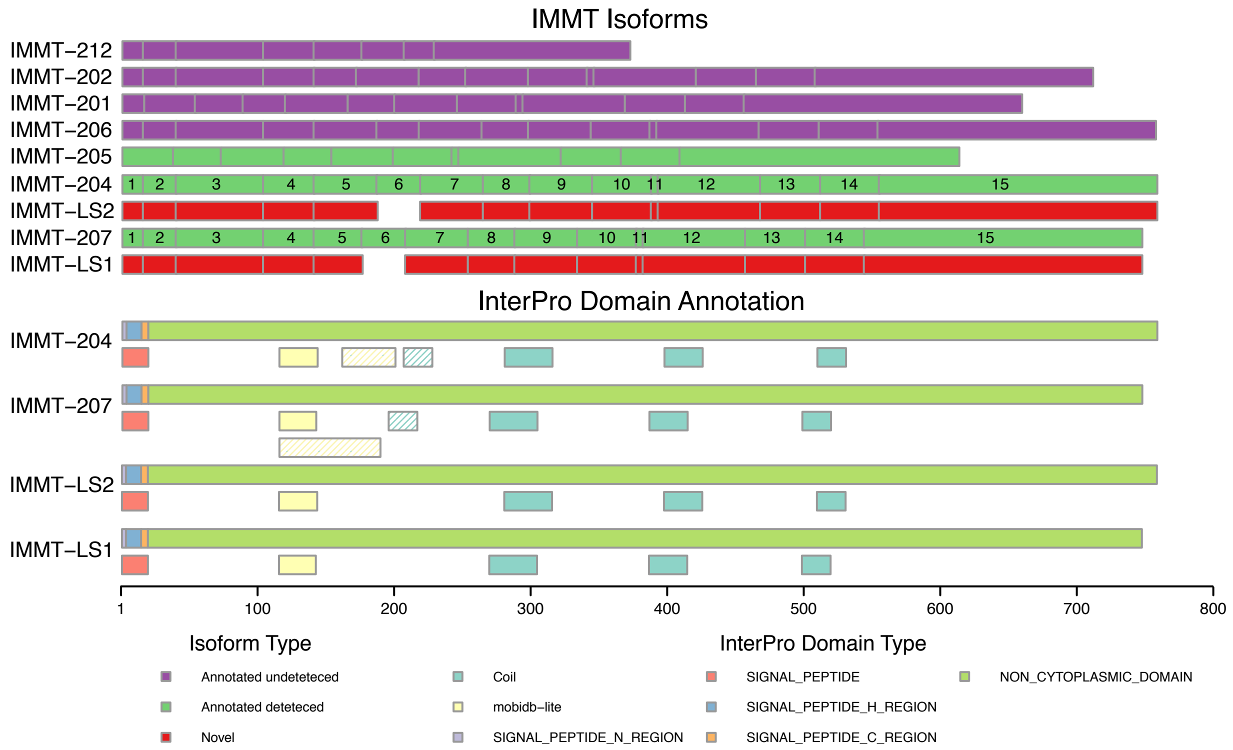


**Supplementary Figure 21. Protein domains of IMMT isoforms annotated by InterPro database.** Exon 6 in IMMT-204 and IMMT-207 encodes 32 amino acids which is a part of a coil-coil and an intrinsic disordered domain (striped) based on annotation from the InterPro database. The absence of exon 6 in the novel transcript isoforms, IMMT-LS2 and IMMT-LS1 (counterparts for IMMT-204 and IMMT207, respectively) disrupts these functional domains based on the InterPro prediction, presumably resulting in proteins with a different function. IMMT-LS1 and IMMT-LS2 isoforms are aligned to their counterparts for illustration purposes.

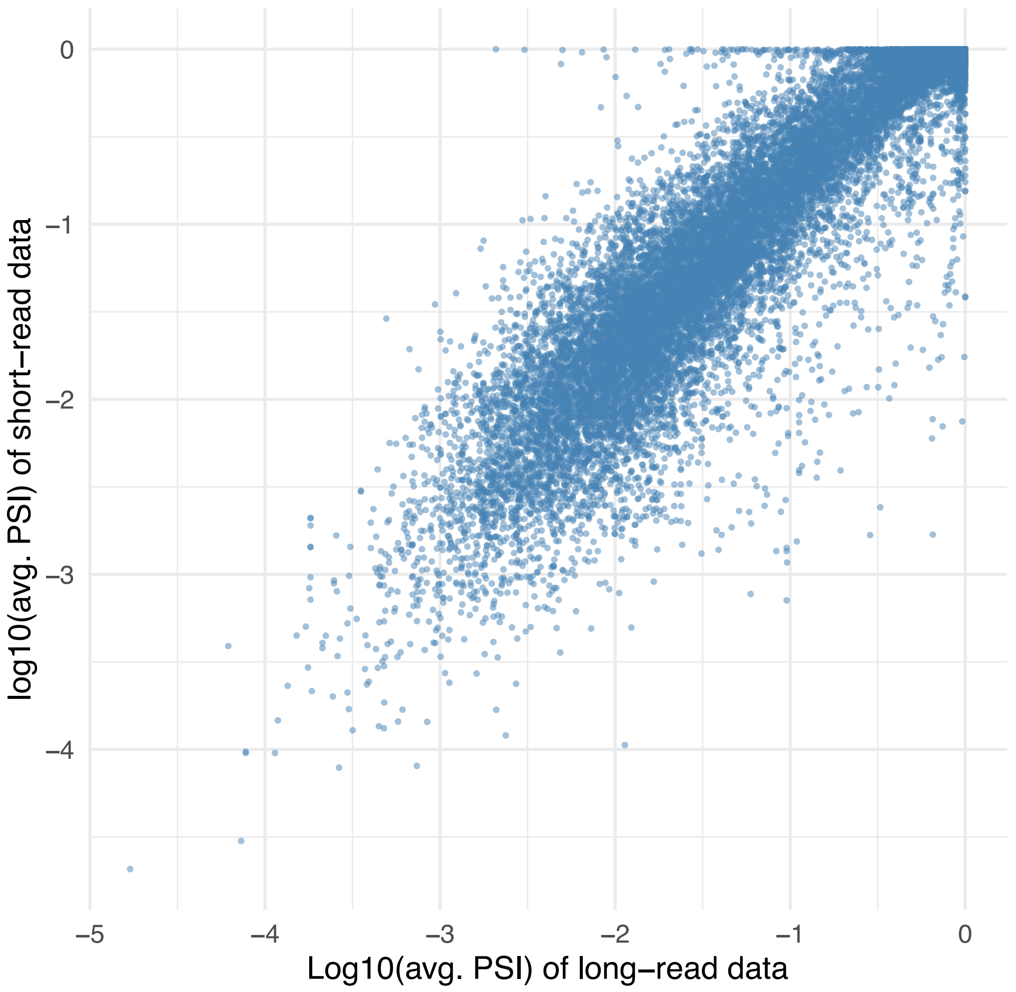


**Supplementary Figure 22. PSI quantification from long and short-read data.** Both datasets show highly comparable PSI values (rank correlation coefficient of 0.7) for exons of interest. PSI is defined as the ratio of number of inclusive reads over total number reads for a exon of interest. Shown are log10 transformed PSI values averaged (median) across all samples. Only exons with long-read coverage over 10 and short-read coverage 100 are included in the plot.

#### List of Oligonucleotides

1. Guide RNA cloning, with lower-case letters represent overhang nucleotides for cloning
Fw_Guide2: CACCGCTCACCGGACTACGAGACCG
Rv_Guide2: aaacCGGTCTCGTAGTCCGGTGAGC

2. Single-stranded donor DNA, with the lower-case letter represents the R634Q point mutation:
TGTGGGACCTCGGGGAGAGTGACCGGCTCACCGGACTACGAGACtGCGGCCTTTC
TGGGCCATATCTGTGAGGGAGCCAAGGAGCAGGATTTAGAATCTTCACACCTCCC
ATCCCACCCCACCCACA.

3. PCR amplification for target *RBM20* genomic region:
RBM20_Fw: CTGGACTAGGGCAATCTTGCCC
RBM20_Rev: CTCATTCTGCTTGGCCTTGGCG

#### Supplementary Tables

Explanation of the columns (in case not self-explanatory) are included as comments (marked with #) above the table.

**Supplementary Table 1. ONT sequencing runs with yield for all samples.**

**Supplementary Table 2. All identified transcripts isoforms with exon coordinates in BED format.**
**Supplementary Table 3. Quantification (read counts) of all transcript isoforms in all samples.**
**Supplementary Table 4. Significantly co-associated exon pairs.**

**Supplementary Table 5. List of differentially expressed transcript isoforms between wildtype and mutants.**

**Supplementary Table 6. List of oligos for validating novel splice events with fragment analysis and validation status.**
